# Hyocholic acid species and the risk of type 2 diabetes

**DOI:** 10.1101/503532

**Authors:** Xiaojiao Zheng, Tianlu Chen, Runqiu Jiang, Aihua Zhao, Fengjie Huang, Yunjing Zhang, Xiaolong Han, Mengci Li, Meilin Wei, Yijun You, Shouli Wang, Xiaojing Ma, Yuqian Bao, Miao Jiang, Jiajian Liu, Qing Zhao, Kun Ge, Bing Dong, Defa Li, Dandan Liang, Sha Lei, Yitao Li, Ke Lan, Aiping Lu, Weituo Zhang, Congrong Wang, Haoyong Yu, Cynthia Rajani, Jun Panee, Guoxiang Xie, Weiping Jia, Wei Jia

## Abstract

Hyocholic acid (HCA) and its derivatives are found in only trace amounts in human blood, but constitute approximately 76 % of the bile acid (BA) pool in the pig, a species known for its exceptional resistance to type 2 diabetes mellitus (T2DM). Here we show that HCA species play a crucial role in maintaining glucose homeostasis and preventing T2DM. We found that in two cohort studies (n=1,213), both obesity and diabetes were associated with lower serum concentrations of HCA species. Serum HCA levels in apparently healthy individuals (n=132) were found to be strong predictors for metabolic health 10 years later. Oral administration of HCA increased serum fasting GLP-1, to a greater extent than metformin, in healthy and diabetic mouse models. HCA upregulated GLP-1 secretion in intestinal enteroendocrine cells via simultaneously activating G-protein-coupled BA receptor, TGR5, and inhibiting farnesoid X receptor, a unique mechanism that is not found in other BA species.

## INTRODUCTION

Bile acids (BAs) have long been regarded as digestive detergents for cholesterol elimination, but are emerging as important signaling molecules that regulate the metabolism of triglyceride, cholesterol, and glucose ^1,2^, and thus, are critically involved in the development of type 2 diabetes mellitus ^3,4^. Glucagon-like peptide-1 (GLP-1) is an incretin hormone that enhances insulin secretion and decreases blood glucose. The expression and secretion of GLP-1 in enteroendocrine L-cells is regulated by two BA receptors, i.e., cell membrane G-protein-coupled BA receptor TGR5 ^5,6^ and nuclear farnesoid X receptor (FXR) ^7^, suggesting that BAs and BA analogs may be used to improve glucose homoeostasis. In support of this view, altered BA profiles were found in patients who underwent bariatric surgery for weight and T2DM control ^8^. Increases in the BA pool size and individual BA species occurred rapidly after the surgery, even before there was significant weight loss ^9,10^.

The composition of the BA profile varies markedly among mammalian species. A recent study reported that hyocholic acid (HCA, also known as 3α,6α,7α-trihydroxy-5β-cholanic acid, and gamma-muricholate) and its glycine- and taurine-conjugated derivatives constituted ∼42 % of total BAs in pig plasma, but comprised only ∼1 % in the plasma of human and rat ^11^. Pigs are routinely raised on obesogenic diets and have little physical activity, which represent a typical diabetogenic condition for humans. However, pigs are resistant to the spontaneous development of T2DM, even after induction with high fat, high fructose and high carbohydrate diets ^12,13^. Because of this metabolic feature, pigs have been used to study hypoglycemia ^14^. We suspected that the distinct BA profile, i.e., the high abundance of HCA and its derivatives in pigs, may play a role in regulating glucose homeostasis leading to their exceptional resistance to metabolic disorders.

To test this hypothesis, we measured the concentrations of HCA species in the serum and feces of diabetic patients and healthy controls and evaluated the predictive value of HCA species for future metabolic outcome for patients. We then validated the effect of HCA species in three mouse models and one pig model. Finally, we assessed the effects of HCA species on GLP-1 expression and secretion in intestinal enteroendocrine L-cells, and the roles of TGR5 and FXR in HCA species-mediated GLP-1 upregulation. This study underscores a critical role of HCA species in maintaining glucose homeostasis in human and other mammalian species, and suggests potential pharmaceutical applications of this group of BAs.

## RESULTS

### Lower levels of serum HCAs in diabetes

To evaluate the association between HCA species and diabetes, we conducted a targeted serum BA profiling in a cohort consisting of 1,107 participants (610 men and 497 women) selected from the Shanghai Obesity Study ^15^. The participants were separated into three groups: healthy lean (HL, n=585), healthy overweight/obese (HO, n=419), and overweight/obese with newly diagnosed T2DM (OD, n=103). Key clinical metabolic markers were significantly different between any 2 of the 3 groups (Table S1). Although the 3 groups had similar total BA (TBA) levels in all, men, and women groups (Fig S1), total concentration of HCA species, i.e. the concentration summation of HCA, hyodeoxycholic acid (HDCA), glycohyodeoxycholic acid (GHDCA), and glycohyocholic acid (GHCA), was the highest in HL and lowest in OD (Fig S1). In addition, the HCA, HDCA, GHDCA, and GHCA concentrations (Figs S1, S2) were decreased in HO and even more so, in OD relative to HL. Pairwise Spearman correlation analysis showed that total and individual HCA species inversely correlated with fasting and post-load glucose, insulin levels and insulin resistance shown by HOMA-IR (Fig S3).

From HL to HO to OD, the participants had increasingly older age, higher body mass index (BMI), and a lower ratio of men/women (although the sex ratios were not significantly different among groups) (Table S1). To eliminate the confounding effects of age, sex, and BMI, we selected 103 older participants with higher BMI, and more women from the HL and HO groups to better match the 103 participants in the OD group. After this selection, all 3 groups had matched age and sex ratios while HO and OD also had matched BMI (Table S2). The 3 groups had similar TBA levels (Fig 1a) and gradually decreased levels of total (Fig 1b) and individual HCA species (Figs 1c, 1k-1n) after this selection, with the fold changes of HO/HL and OD/HL for total HCA species (0.75 and 0.55, respectively), HCA (0.82 and 0.45), HDCA (0.81 and 0.47), GHCA (0.68 and 0.57), and GHDCA (0.72 and 0.60). HCA species remained inversely correlated with fasting and post-load levels of glucose and insulin as well as, insulin resistance after the selection (Figs 1d-1j). The results suggest that obesity (HO+OD vs. HL) and diabetes (OD vs. HO) were associated with lower concentrations of total and individual HCA species in serum.

**Figure 1.**
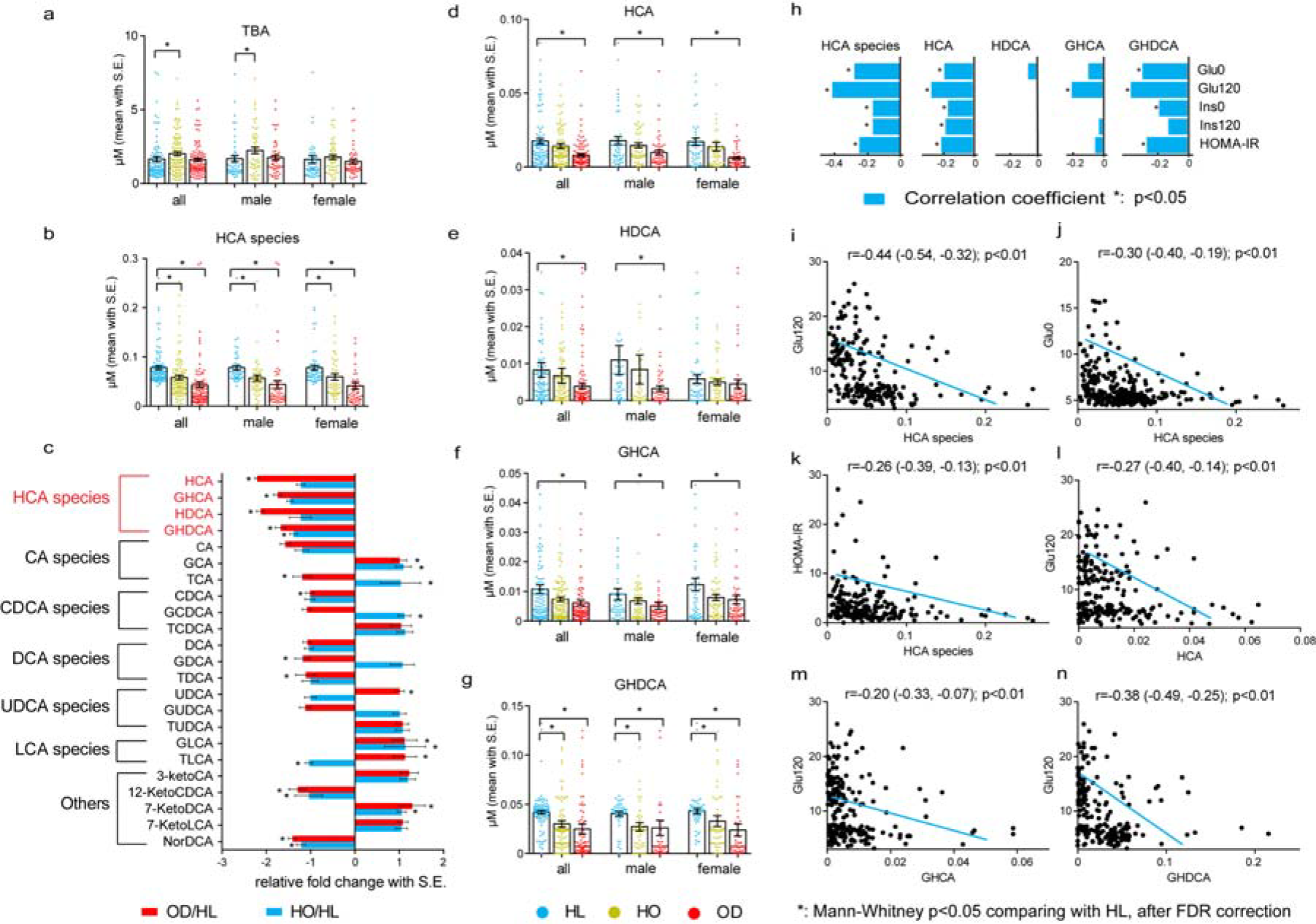
Performances of HCA species in the first cross-sectional study. (a) Total bile acid (TBA) and (b) HCA species levels (mean with S.E.) in matched healthy lean (HL, n=103 from 585), healthy overweight/obese (HO) (n=103 from 419) and overweight/obese with type 2 diabetes (OD) (n=103) groups. * Corrected (FDR=0.05) Mann-Whitney p<0.05 when compared with HL. (c) Fold of change (mean with S.E.) of 23 BAs in HO and OD groups relative to HL group. * FDR corrected Mann-Whitney p<0.05 when compared with HL. Levels of HCA species (HCA, HDCA, GHCA and GHDCA, highlighted in red) were consistently lower in HO and OD groups compared with HL group. (d -g) Group differences (mean with S.E.) of individual HCA species based on matched all (n=309), male (n=156) and female (n=153) samples. * FDR corrected Mann-Whitney p<0.05 when compared with HL. (h) Correlation coefficients of total and individual HCA species with representative metabolic markers (matched samples). * p < 0.05. (i -n) Scatter plots of total or individual HCA species versus representative metabolic markers.

### Development of diabetes associated with depleted HCA levels

To confirm the findings above in a separate cohort, and to evaluate the association between fecal HCA and diabetes, we recruited a second cohort of 106 participants (44 men and 62 women), which included 32 healthy, 34 pre-diabetic and 40 diabetic individuals. The HbA1c and fasting and post-load blood glucose levels of pre-diabetic and diabetic patients were significantly higher than those of healthy controls (Tables S3, 4). No significant group differences were found in serum and fecal total BAs (Figs 2a, c). Compared with healthy controls, the pre-diabetic and diabetic groups had lower levels of total HCA species in both serum and feces, in the groups, all, men, and women (Figs 2b, d and Tables S5, 6). The group differences were greater in feces than in serum. As expected, individual HCA species showed similar group differences (Figs 2e-j). The concentrations of fecal GHCA and GHDCA are not shown as they were below the detection limit. Total and individual HCA species in feces had stronger inverse correlations with fasting and post-load blood glucose levels than serum levels of HCA species (Figs 2k-m, adjusted for age, sex and BMI).

**Figure 2.**
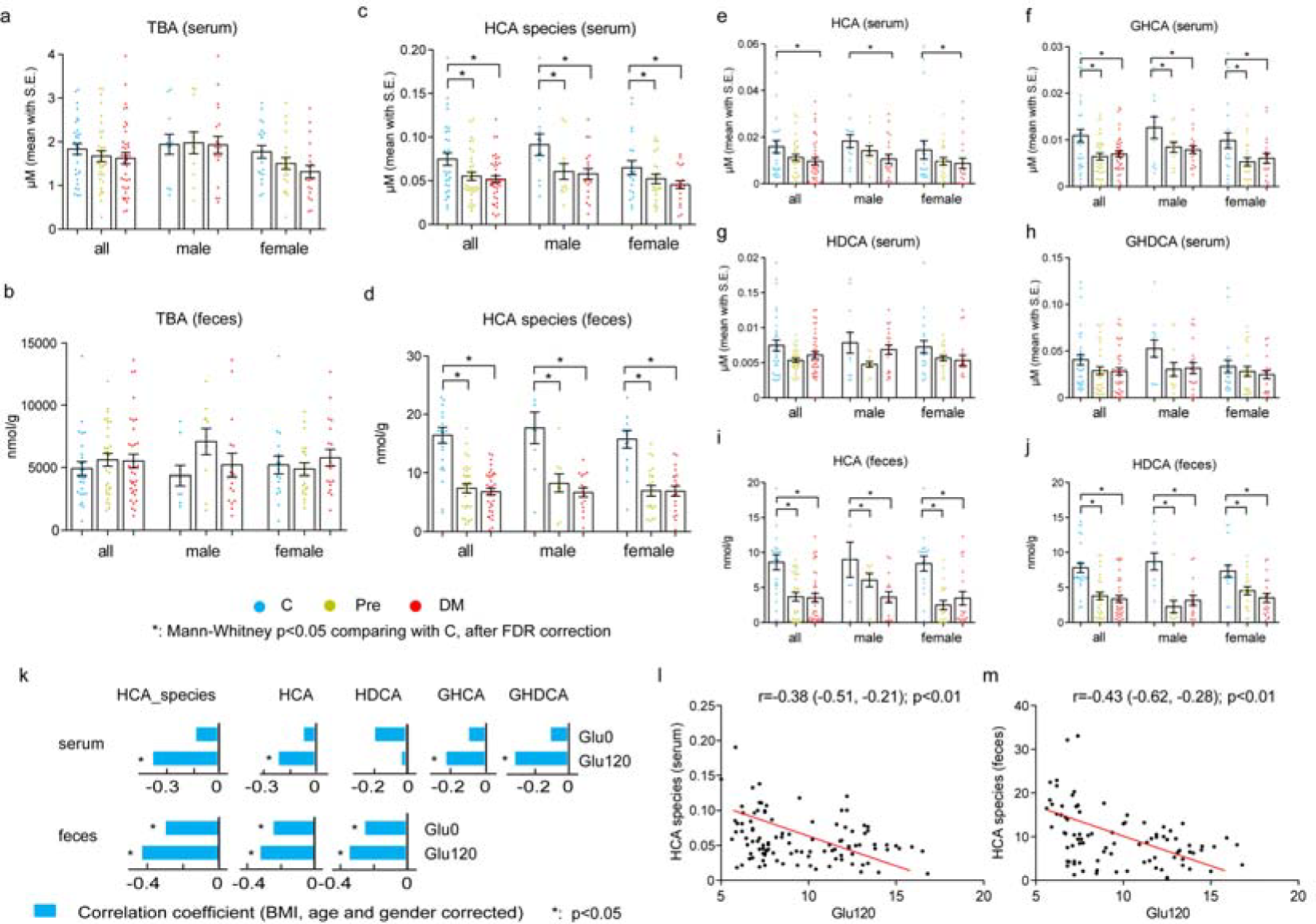
Performance of HCA species in the second cross-sectional study. (a -d) Total bile acids (TBA) and total HCA species in serum and feces in healthy control (C, n=32), pre-diabetes (Pre, n=34) and diabetes (DM, n=40) groups. (e -j) Individual HCA species in the 3 groups in all (n=106), male (n=44) and female (n=62) samples. Mean with S.E. * FDR corrected Mann-Whitney p<0.05 when compared with C. (k) Correlation coefficients of total and individual HCA species in serum and feces with glycemic markers. * p<0.05, adjusted for BMI, age and sex. (l, m) Scatter plots of total HCA species in serum or feces versus a representative glycemic marker.

### HCAs were predictors for metabolic outcome

To evaluate the association between HCA species and future metabolic health, we selected 132 subjects (36 men and 96 women) from the Shanghai Diabetes Study ^16^. All of them were metabolic healthy (MH, defined in the Method Section) at their enrollment. After 10 years, 86 participants became metabolically unhealthy (MU, defined in the Method Section), and 46 remained MH. At baseline, the future MU group were older, had higher BMI and more men than the future MH group (although group differences of sex ratio did not reach statistical significance), however, the major metabolic markers were similar between the two groups (Table S7). To eliminate the confounding effects of age, sex and BMI, we chose 46 younger participants with lower BMI and comprised of more women, from the MU group to match the 46 participants in the MH group (Table S8). When samples from all participants were considered, the concentrations of total BAs in serum were comparable between the MH and MU groups, but the concentrations of total and individual HCA species were significantly lower in the MU than the MH group (Fig S4 and Table S9). Age-, sex- and BMI-matched samples yielded similar results as all samples did (Figs 3a-f, Table S10), suggesting that the baseline differences of HCA species between MH and MU groups were independent of age, sex, and BMI. Binary logistic regression analysis of all samples showed that the association between HCA species and future MU outcome were (odds ratio (95 % CI) 0.89 (0.86, 0.93), 0.91 (0.87, 0.94), 0.90 (0.84, 0.96), 0.92 (0.85, 0.99), 0.52 (0.40, 0.69) and 0.90 (0.86, 0.94)) for total HCA species, HCA, GHCA, HDCA, and GHDCA, respectively (p<0.05 for all, adjusted for age, sex, and BMI) (Fig S5). The receiver operating characteristic (ROC) curve analysis showed that total HCA species (red line in Fig 3g) had the highest area under curve (AUC) of 0.83, and the AUCs of individual HCA species ranged from 0.63 to 0.79, providing supporting evidence for using total and individual HCA species as predictors for future metabolic outcome.

**Figure 3.**
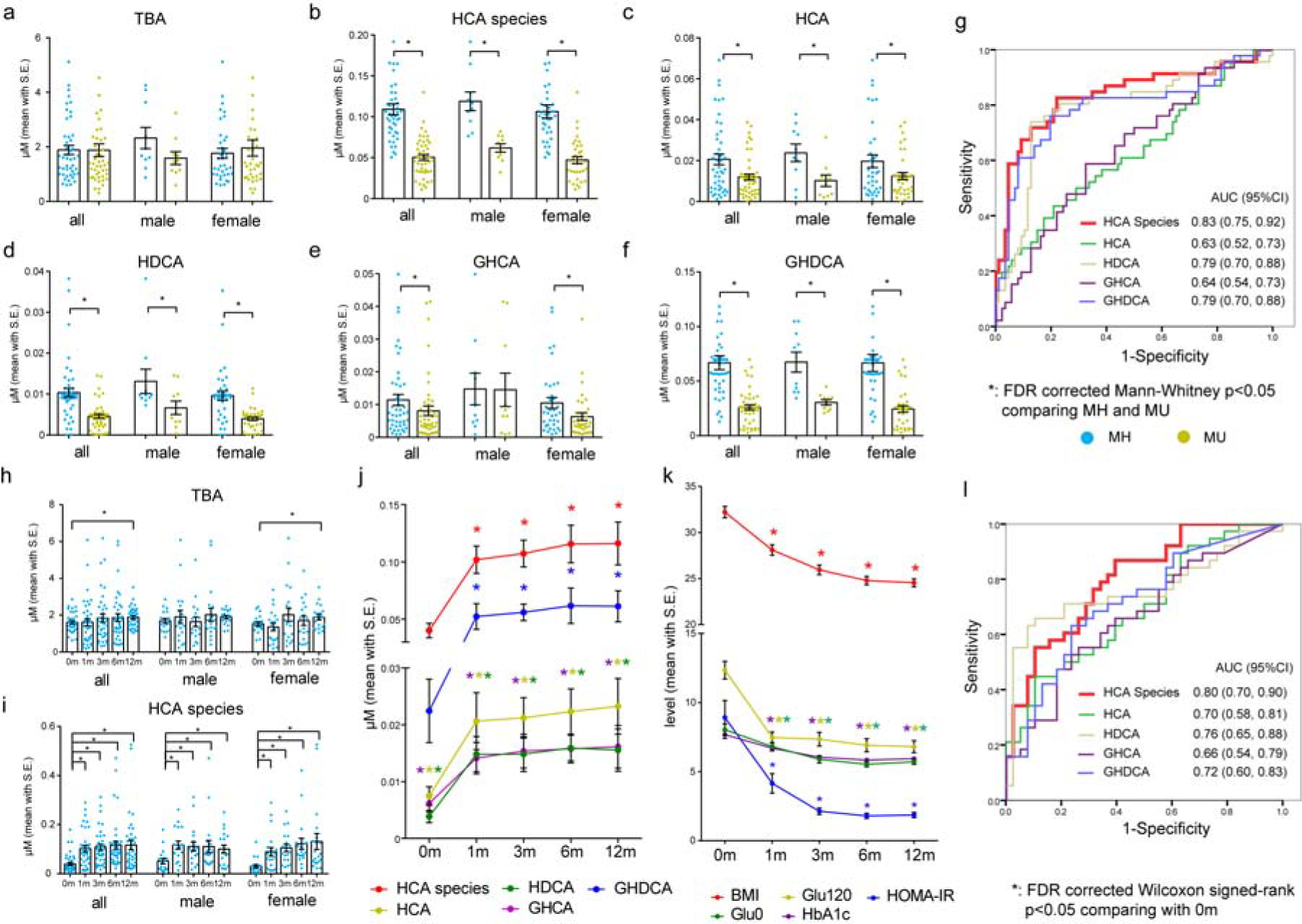
Performances of HCA species in the 10-year longitudinal study and surgery-induced changes of HCA species in the gastric bypass surgery intervention study. (a -f) Total bile acids (TBA), total and individual HCA species in serum of age and BMI matched all (n=92), male (n=20) and female (n=72) individuals in future metabolically healthy (MH) and metabolically unhealthy (MU) groups. Mean with S.E., * FDR corrected Mann-Whitney p<0.05 when comparing MH and MU. (g) Receiver operating characteristic (ROC) analyses of total and individual HCA species for the metabolic health longitudinal study (all samples). (h, i) TBA and total HCA species serum concentrations before and after gastric bypass surgery in 38 obese and diabetic patients (j) Serum concentrations of total and individual HCA species before and after surgery (k) BMI and glycemic markers before and after surgery. * FDR corrected Wilcoxon signed-rank test p<0.05 when compared with baseline (0m). (l) ROC analysis of the changes (12 months vs. baseline) of total and individual HCA species.

### Gastric bypass surgery increased serum HCAs

We further studied the changes of HCA species in diabetic patients after Roux-en-Y gastric bypass (RYGB) surgery. Thirty-eight obese diabetic patients who received RYGB were examined before and at 1, 3, 6, and 12 months post-surgery (Table S11). Serum concentrations of total BAs gradually increased after RYGB surgery, and became significantly higher than baseline at 12 months post-operation (Fig 3h). The concentrations of total and individual HCAs in the serum increased drastically 1 month after the surgery (FC = 2.52, 2.75, 3.86, 2.28, and 2.33 for total HCA species, HCA, HDCA, GHCA, and GHDCA, respectively) and maintained minor increases afterwards (Figs 3i-j, Table S12). Improvements of BMI, fasting and post-load blood glucose levels, HbA1c, and insulin resistance occurred throughout the 12 months (Fig 3k). ROC analysis showed that the AUCs of the 12-month changes of total HCA species, HCA, GHCA, HDCA, and GHDCA were 0.80, 0.70, 0.66, 0.76, and 0.72, respectively (Fig 3l), evidence for potential prediction capability for the metabolic outcome of RYGB surgery.

### HCAs regulated blood glucose and GLP-1 in animal models

To understand the potential role of HCA species in regulating glucose homeostasis in pigs, we compared the BA profiles in the sera of pig (n=6, 3 males and 3 females), human (from the first cohort study, n=1,107, 610 men and 497 women), and mouse (wildtype C57BL/6J, n=10, 5 males and 5 females). Fig 4a shows that the HCA species accounted for the majority of BAs in the serum of pig (75.96 ± 4.00 %), but for only very small portions in those of human (4.99 ± 0.14 %) and mouse (3.11 ± 0.12 %). All 6 HCA species, i.e., HCA, GHCA, tauro-HCA (THCA), HDCA, GHDCA and tauro-HDCA (THDCA), were detected in pig serum, but only some of these HCA species were detected in human and mouse. Meanwhile, the fasting blood glucose level was the lowest in pig (4.4 ± 0.1 mmol/L), followed by human (5.5 ± 0.0 mmol/L) and mouse (5.3 ± 0.2 mmol/L) (Fig 4b), which was in the opposite order of the abundance of serum HCA species in these species.

**Figure 4.**
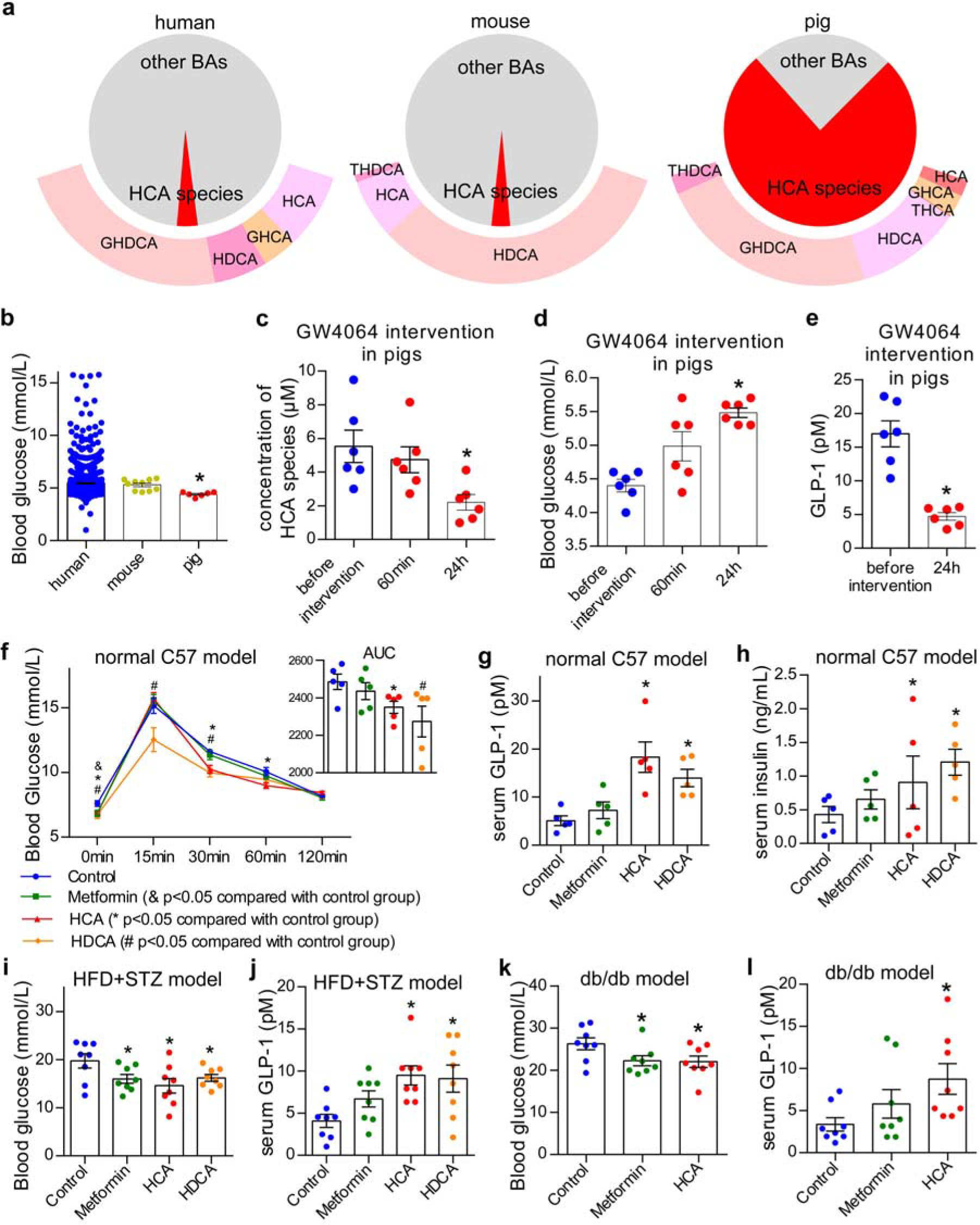
Effects of HCA species on the levels of blood glucose, GLP-1 and insulin in animal models. (a) BA composition in the serum of humans, C57BL/6J mice, and pigs. The pie charts are composed of HCA species (red) and other BAs (grey). The outer rings are composed of detected individual HCA species. (b) The fasting blood glucose levels of humans, mice, and pigs. (c) Serum concentrations of total HCA species and (d) blood glucose before and 60 min and 24h after GW4064 oral administration (10 mg/kg, twice with a 12 h interval) in pigs. (e) Serum GLP-1 level before and 24 h after GW4064 treatment in pigs. (f) Blood glucose levels and AUC of OGTT, (g) serum GLP-1 levels, and (h) insulin levels of normal C57BL/6J mouse models treated with metformin (200 mg/kg/day), HCA (100 mg/kg/day), HDCA (100 mg/kg/day) and vehicle control for four weeks. (i) Blood glucose levels, and (j) serum GLP-1 levels of HFD+STZ mouse models treated with metformin (200 mg/kg/day), HCA (100 mg/kg/day), HDCA (100 mg/kg/day) and vehicle control for four weeks. (k) Blood glucose levels and (l) serum GLP-1 levels (g) of db/db mouse models treated with metformin (200 mg/kg/day), HCA (100 mg/kg/day) and vehicle control for four weeks. Mean with S.E. * p<0.05 when compared with control group using unpaired t-test, except compared with before intervention group in pig model using paired t-test.

We further treated the pigs with GW4064, a FXR agonist, via oral gavage at a dose of 10 mg/kg (twice with a 12 h interval), in an effort to suppress hepatic BA synthesis. This was done to answer the question whether GW4064 would reduce serum HCA levels in pigs and furthermore, whether HCA depletion would decrease circulatory GLP-1 concentration and increase blood glucose levels. After GW4064 treatment, the concentration of HCA species in serum decreased by 60 % (Fig 4c, and Figs S6a-g). Meantime, the blood glucose levels increased by 25 % (Fig 4d) and that of serum GLP-1 decreased by 72 % (Fig 4e). Blood glucose levels were also measured 15 and 35 minutes after GW4064 treatment, the data and interpretation can be found in Figs S6h-i.

To investigate whether HCA species have direct impact on glucose homeostasis, we treated healthy C57BL/6J mice for 4 weeks with HCA (100 mg/kg/day), HDCA (100 mg/kg/day), metformin (200 mg/kg/day), and 6 % sodium bicarbonate (NaHCO_3_) as vehicle control. Mice in metformin, HCA, and HDCA groups showed improved oral glucose tolerance at 4 weeks (Fig 4f). The hypoglycemic effect was more rapid with HCA species intervention (significant at 1 week) compared to metformin (significant at 4 weeks) (Figs S7a-d). Moreover, mice treated with HCA and HDCA showed higher circulating GLP-1 levels (Fig 4g) and fasting insulin (Fig 4h) than metformin at 4 weeks.

We then investigated whether HCA could improve glucose homeostasis under obese and diabetic conditions in a high-fat diet-streptozotocin (HFD + STZ) induced diabetic and a db/db mouse model. For the HFD + STZ model, mice were treated with HCA (100 mg/kg/day), HDCA (100 mg/kg/day), metformin (200 mg/kg/day), and 6 % NaHCO_3_ as vehicle control, respectively. At 4 weeks, mice treated with metformin, HCA, or HDCA showed significantly lower fasting blood glucose levels than controls (Fig 4i). Similarly, the hypoglycemic effect was more rapid with HCA or HDCA treatment compared to metformin (Figs S7e-h). Furthermore, mice treated with HCA or HDCA showed increased circulating GLP-1 levels (Fig 4j). In a db/db mouse model, mice were treated with HCA (100 mg/kg/day), metformin (200 mg/kg/day), and vehicle control. At 4 weeks, db/db mice showed significantly lower fasting blood glucose levels in metformin and HCA treatment groups (Fig 4k, Figs S7j-m), higher circulating GLP-1 levels (Fig 4l) and higher fasting insulin in HCA group (Fig S7n), compared to controls.

### HCAs upregulated GLP-1 via TGR5 and FXR signaling

We compared the responses of intestinal enteroendocrine STC-1 and NCI-H716 cells ^17,18^ to HCA species and other BAs on GLP-1 transcription and protein expression. The results showed that no apparent GLP-1 upregulation using all BAs at 5 μM (Figs S8a, b). When the concentration increased to 25 μM, all of the BAs upregulated GLP-1 transcription and protein expression (Figs S8c-e), among which, HCA species were most effective. At 50 μM, HCA species upregulated GLP-1 transcription and protein expression significantly more than HCA species at 25 μM (Figs 5a, b), while other BAs did not upregulate GLP-1 expression. These results showed the difference between HCA species and other BAs on regulating the GLP-1 expression, in that the effect of GLP-1 stimulation was dose dependent with HCA species, while the effect was suppressed with other BAs at relatively high concentrations.

**Figure 5.**
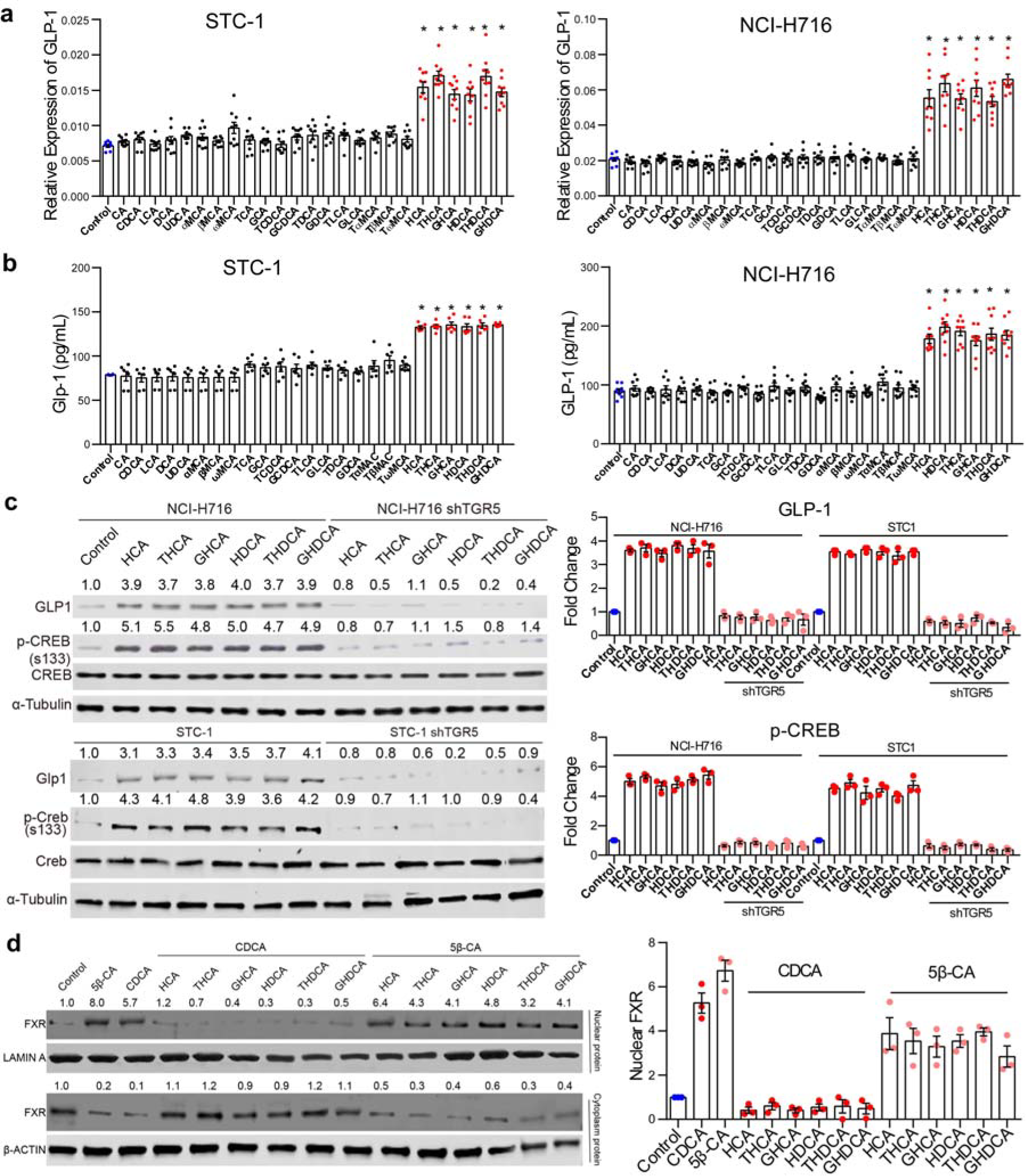
HCA species more effectively upregulated GLP-1 protein expression in enteroendocrine cell lines than other bile acids through effects of TGR5 and FXR. NCIH716 and STC-1 cells were treated with all 6 HCA species and 19 other BAs, each at 50 μM for 48 h. (a) The GLP-1 transcription was measured using Real-time PCR. (b) The GLP-1 secretion was measured using ELISA. (c) NCI-H716 and STC-1 as well as their TGR5 knockdown cells were treated with 6 HCA species for 24h, and intracellular GLP-1, p-CREB and total CREB were determined using western blot. (d) FXR protein concentration in nuclear and cytosolic fractions of NCI-H716 cells treated with 50 μM of CDCA or 5β-CA for 24 hours, with or without the presence of HCA species, each at 50 μM. Representative images are shown, and data were obtained from 3 independent experiments. Mean with S.E. * p<0.05 when compared with control using unparied t-test.

Two BA receptors, TGR5 and FXR, are involved in regulating the GLP-1 expression in enteroendocrine L-cells. We found that each HCA species significantly increased the level of GLP-1 secretion as well as CREB phosphorylation (S133) (p-CREB) (a marker of TGR5 activation) (Fig 5c, left panel of western-blot and bar chart; Figs S9a, b), compared to other BAs. However, GLP-1 and p-CREB expression levels were significantly decreased in TGR5 knockdown cells (Fig 5c, right panel of western-blot and bar chart, Fig S10a,b), suggesting that the upregulation of GLP-1 by HCA species was TGR5 dependent.

Our results showed that two FXR agonists, chenodeoxycholic acid (CDCA) and 5β-Cholanic acid (5β-CA)^19,20^ increased nuclear translocation (a marker of FXR activation) of FXR, and such effect was inhibited by the co-treatment of HCA species (Fig 5d). Western-blot analysis of FXR translocation and SHP expression, one of the downstream proteins of FXR activation, verified the inhibitory effect of HCA species on FXR (Figs S9a, b). Interestingly, non-HCA BAs, at 25 μM, promoted GLP-1 expression via TGR5 activation while their FXR binding and activation was not strong. At higher concentrations (50 μM), there was marked upregulation of FXR by non-HCA BAs (Figs S9a, b) but the GLP-1 production was suppressed.. Such observation was further verified in FXR knockdown cells, where GLP-1 transcription and protein expression was increased significantly with non-HCA BAs intervention in shFXR cells compared to control due to the loss of FXR. No obvious difference was observed between HCA and non-HCA treatments (Figs S11b-d). Previous studies have identified 5β-CA as both a FXR agonist and a TGR5 antagonist, and as expected, the upregulation of GLP-1 by HCA was abolished by 5β-CA co-treatment as shown by transcription (Fig S12a), ELISA (Fig S12b), western blot (Fig S12c), and 2D and 3D IF staining (Figs S12d,e).

We also intended to understand whether the inhibition of FXR by HCA species directly regulated GLP-1 secretion independent of TGR5 signaling, or such inhibition also regulated TGR5 expression and subsequently regulated GLP-1 expression. The control and shFXR cells (Fig S11a) were exposed to HCA species and 4 other representative BAs, cholic acid (CA), CDCA, LCA, deoxycholic acid (DCA). FXR knockdown had no apparent effect on TGR5 and p-CREB expression (Fig S13), suggesting that the effect of TGR5 expression and activation by HCA species was not regulated by FXR. Taken together, in enteroendocrine L-cells, BAs induce GLP-1 secretion through BA- TGR5 and FXR signaling. More specifically, BA-TGR5 signaling promotes GLP-1 expression, whereas BA-FXR signaling inhibits GLP-1 expression. HCA species promoted GLP-1 expression and secretion through a unique mechanism that involved both action as an agonist for TGR5 and action as an antagonist for FXR, simultaneously.

To validate whether HCA species induced GLP-1 secretion depended on TGR5 activation as well as FXR inhibition, we conducted *in vivo* studies for 4 weeks using 5β-CA (100 mg/kg/day, i.g.) to inhibit TGR5 and activate FXR simultaneously, as well as Fexaramine (FEX; 100 mg/kg/day, i.g.) to activate only intestinal FXR. The results (Fig 6a) showed that 5β-CA intervention significantly inhibited HCA-induced GLP-1 secretion. Such inhibition was not as strong with FEX treatment as with 5β-CA treatment, because the presence of HCA-TGR5 signaling was still significant. Meanwhile, HCA induced insulin secretion and blood glucose reduction was reversed by 5β-CA, but was attenuated, to some extent, by FEX (Figs 6b, c).

**Figure 6.**
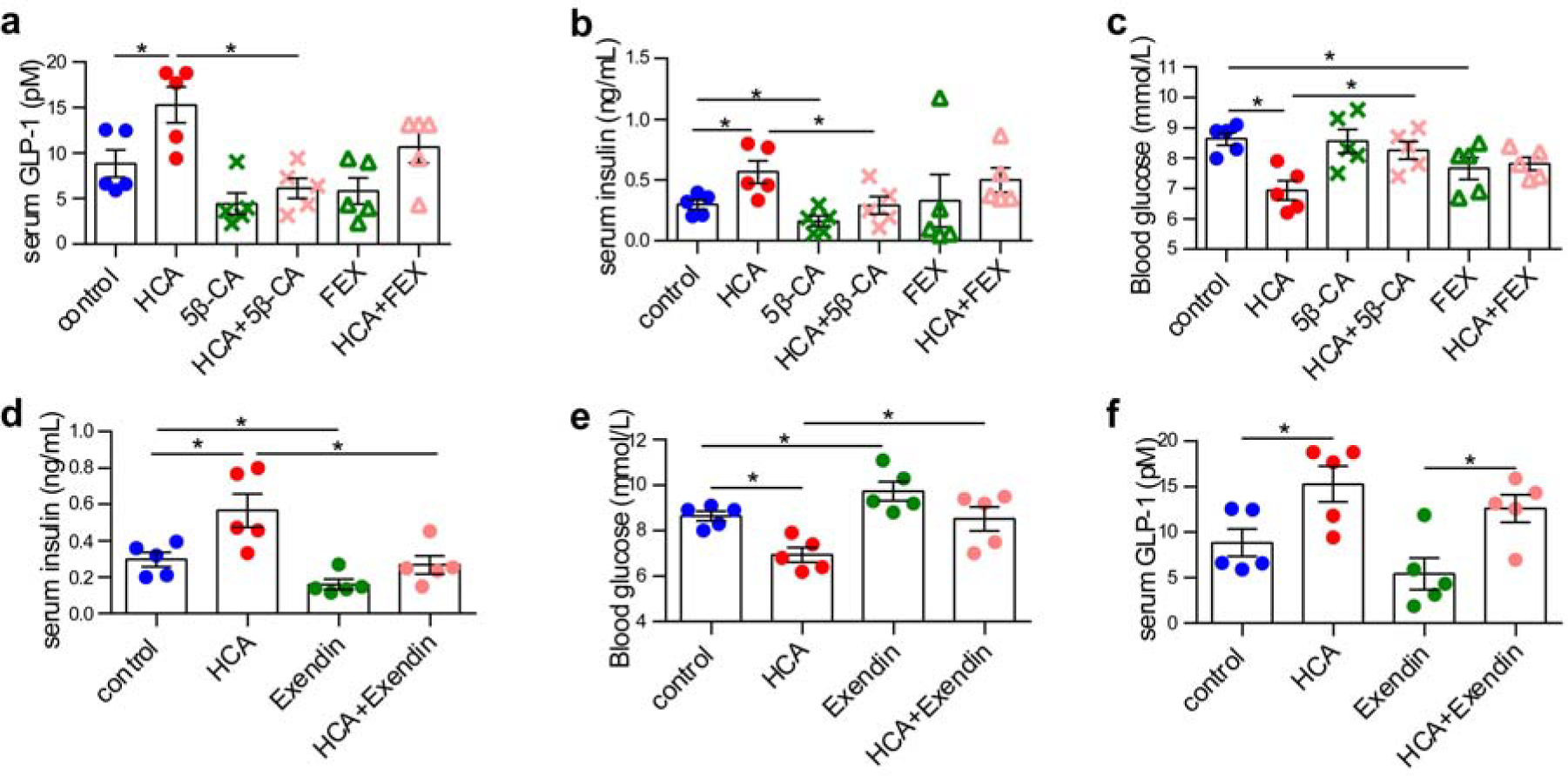
Effect of HCA on the levels of GLP-1, insulin, and blood glucose with TGR5, FXR and GLP-1 receptor intervention. The levels of (a) serum GLP-1, (b) serum insulin, and (c) blood glucose of normal C57BL/6J mice in control, HCA (100 mg/kg/day, i.g.), 5β-CA (TGR5 antagonist and FXR agonist; 100 mg/kg/day, i.g.), HCA+5β-CA, FEX (FXR agonist; 100 mg/kg/day, i.g.), and HCA+FEX groups at 4 weeks. The serum levels at 4 weeks of (d) insulin, (e) glucose, and (f) GLP-1 of normal C57BL/6J mice in control, HCA, Exendin-3(9-39) amide (Exendin, GLP-1 receptor antagonist; 25 nmol/kg/day, i.p.) and HCA+Exendin groups. Mean with S.E. * p<0.05 when compared with control group using unpaired t-test.

We further determined whether HCA induced GLP-1 secretion was an essential pathway involved in HCA regulated glucose metabolism. We inhibited the GLP-1 receptor in a mouse model using a GLP-1 receptor antagonist, Exendin-3(9-39) amide (Exendin; 25 nmol/kg/day, i.p.) for 4 weeks, HCA induced insulin secretion and hypoglycemic effects were abolished (Figs 6d,e).

## DISCUSSION

Among the HCA species, HCA and HDCA were first isolated by Windaus from pig bile ^22,23^. The biosynthetic pathways and physiological levels of HCA and HDCA are different among mammalian species. Synthesis of HCA and HDCA in humans is not fully understood. Early *in vitro* data demonstrated that HDCA can be synthesized from TLCA and LCA via 6α-hydroxylation in human liver microsomes ^24,25^. This pathway was later confirmed ^26^ and attributed to the function of CYP3A4 ^27,28^. A study from the same group reported that HCA can also be synthesized from CDCA through the same CYP3A4-mediated 6α-hydroxylation pathway ^1,29^. Furthermore, HCA and HDCA can also be synthesized from CDCA via hepatic CYP3A1 in combination with gut microbial epimerase enzymes. In rats, HDCA can be synthesized via bacterial biotransformation of β-muricholic acid ^30^, or synthesized from LCA by hepatic enzymes that convert LCA to 3α,6β-dihydroxy cholanoic acid that can be further oxidized by gut bacteria to 3α-hydroxy-6-keto cholanoic acid, and then reduced to HDCA ^31^.

As a key incretin, GLP-1 is produced and secreted by the intestinal enteroendocrine cells. Our *in vitro* data showed that HCA species upregulated GLP-1 gene and protein expression and secretion in intestinal enteroendocrine NCI-H716 and STC-1 cells more effectively than other BA species tested. This was achieved through the simultaneous activation of TGR5 and inhibition FXR by unique interactive signaling of HCA species that has not been observed for other BA species. Our animal studies also showed simultaneous changes in GLP-1 and glucose levels in the blood following HCA species treatment. The effect of HCA species on blood glucose regulation was more potent than the antidiabetic agent, metformin. Therefore, the regulatory effect of HCA species on glucose homeostasis is mainly mediated through promotion of intestinal secretion of GLP-1.

An interesting finding in our study was that although all of the BAs including HCA species have the effect on stimulating GLP-1 secretion, the dose effects were different. At lower concentrations (25 μM), all of the BAs promoted GLP-1 secretion. However, HCA species upregulate GLP-1 secretion in a dose-dependent manner, while other BA species failed to upregulate GLP-1 secretion at relatively higher concentrations (50 μM). Such a unique feature of HCA species suggested that HCA and derivatives could be applied with sufficiently high concentrations (pharmacological levels) in maintaining glucose homeostasis, thus having great potential for therapeutic applications.

In clinical studies, T2DM is inherently associated with obesity and aging^32^, so we tried to eliminate the confounding effects of BMI and age when evaluating the role of HCA species in T2DM. By matching age and/or BMI between the groups in comparison, we demonstrated that HCA species had direct correlations with glycemic markers and future metabolic outcome. These results provide evidence that HCA species play critical roles in regulating glucose homeostasis and are protective against the development of T2DM in humans.

We also showed that, compared with healthy controls, pre-diabetic and diabetic patients had ∼27 % lower serum levels of HCA species, but strikingly ∼57 % lower HCA species in feces, although these patients had similar levels of total BAs in feces as controls. Notably the pre-diabetic and diabetic patients had higher BMIs than the healthy controls, which suggest that they may also have altered gut microbiota ^33^. Intestinal microbiota are known to play a critical role in BA metabolism ^34-36^. Obesity and/or diabetes-associated changes in gut microbiota may inhibit the generation or facilitate the metabolism of HCA species, which, in turn, could lead to their depletion in feces. We further showed that fecal HCA species had stronger inverse correlations with glycemic markers than serum HCA species after adjusting for age, sex, and BMI, suggesting that the intestinal track is a critical site for HCA-mediated glycemic regulation.

RYGB surgery is considered a rapid resolution of T2DM. Both HCA and GHCA were found significantly increased after RYGB ^11^. We found that in addition to HCA and GHCA, HDCA and GHDCA were also increased drastically after RYGB; and among all BAs, the increases in HCA species were the most pronounced and consistent (Table S12). Our results further highlighted the critical role of HCA in glucose regulation following bariatric surgery and their predictive value for the post-operation metabolic outcome.

## CONCLUSION

The composition of the BA profile especially HCA species varies markedly among mammalian species. We show in this study that obesity and diabetes were closely associated with significant lower levels of HCA species in serum. Furthermore, the concentrations of HCA species in both serum and feces were closely correlated with glycemic markers and were strong predictors of future metabolic outcome in apparently healthy individuals. HCA species were shown to upregulate the gene transcription, protein expression and secretion of GLP-1 in both intestinal enteroendocrine NCI-H716 and STC-1 cells to a significantly greater extent than other BA species. This action was mediated through simultaneous activation of TGR5 and inhibition of FXR. Taken together, our results provide strong supporting evidence that HCA species are protective against the development of diabetes in mammals and have the potential to be used as a treatment for type 2 diabetes. Future research is warranted to further improve our knowledge on the correlations of HCA and gut microbiota in an effort to identify possible probiotic treatment possibilities.

## METHODS

### Human experiments

#### Human study 1: cross sectional study 1

A total of 1,107 fasting serum samples obtained from 585 healthy lean (329 men and 256 women), 419 healthy overweight/obese (229 men and 190 women) and 103 overweight/obese diabetic (52 men and 51 women) participants were selected from the Shanghai Obesity Study ^15^. Individuals were excluded if they had chronic inflammatory disease, cardiopulmonary, renal or liver disease, active malignancy, or were taking any medication (including weight loss or psychotropic medication).

#### Human study 2: cross sectional study 2

A group of 106 subjects including 32 healthy controls (12 men and 20 women), 34 pre-diabetic individuals (12 men and 22 women) and 40 diabetic patients (20 men and 20 women) were recruited for this study. The exclusion criteria were the same as in human study 1. Fasting sera of all the participants and fecal samples of 91 participants (26 healthy controls, 30 pre-diabetes and 35 diabetic patients) were collected and stored for later analysis.

#### Human study 3: 10-year longitudinal study

A group of 132 subjects (36 men and 96 women) were selected from the Shanghai Diabetes Study, which was intended to assess the prevalence of diabetes and diabetes-associated metabolic disorders in urban Shanghai ^16^. All 132 subjects were metabolically healthy at baseline (year 2000-2001). Ten years later (year 2010-2011), 86 participants (26 men and 60 women) became metabolically unhealthy (future metabolically unhealthy) and 46 (10 men and 36 women) remained healthy (future metabolically healthy). Fasting serum samples of the 132 participants at baseline were collected and stored for future analysis.

#### Human study 4: Gastric bypass surgery intervention study

A total of 38 obese diabetic patients who received Roux-en-Y gastric bypass surgery were enrolled in the study ^10^. Any patient with a history of open abdominal surgery, a serious disease (such as heart or lung insufficiency) that was incompatible with surgery, an acute type 2 diabetes complication, severe alcohol or drug dependency, a mental disorder, type 1 diabetes, secondary diabetes, an unstable psychiatric illness, or who was at a relatively high surgical risk (such as a patient with an active ulcer) was excluded. The fasting serum specimens of these subjects were collected and stored for future analysis before (baseline) and 1, 3, 6, and 12 months after the surgery.

#### Clinical measurements

Fasting and 2 h postprandial plasma glucose and insulin levels, serum lipid profiles (total cholesterol TC, triglyceride TG, high-density lipoprotein-cholesterol HDL, low-density lipoprotein-cholesterol LDL), blood pressure (systolic and diastolic blood pressure SP and DP), waist circumference, BMI, liver and kidney function tests were determined as previously described ^38^.

#### Definitions of lean, overweight/obesity, pre-diabetes, diabetes, metabolically healthy and unhealthy

Individuals with BMI < 25 kg/m^2^ were considered lean and those with BMI ≥ 25 were classified as overweight/obese. Individuals with 6.1 mmol/L ≤ fasting blood glucose < 7.0 mmol/L or 7.8 mmol/L ≤oral glucose tolerance test (OGTT) (2 h) < 11.1 mmol/L were classified as pre-diabetic. Subjects with fasting blood glucose ≥ 7.0 mmol/L and/or OGTT (2 h) ≥ 11.1 mmol/L were classified as diabetic. Subjects were considered “metabolically healthy” if they met all of the following criteria: fasting blood glucose < 6.1 mmol/L, OGTT (2 h) < 7.8 mmol/L and no previous history of diabetes; SBP/DBP <140/90 mmHg and no previous history of high blood pressure; fasting plasma TG < 1.7 mmol/L and fasting plasma HDL ≥ 0.9 mmol/L (men) or ≥ 1.0 mmol/L (women), and no previous history of high cholesterol (TC < 5.18 mmol/L); no history of cardiovascular or endocrine disease ^39^. Those who failed to meet all criteria above were classified as “metabolically unhealthy”.

#### Sample collection

All human samples were collected and stored following the standard operating protocol of the hospital. Briefly, fasting venous blood samples were obtained before 10 AM and were centrifuged immediately. The serum was removed from the cells, divided into aliquots and delivered on dry ice to the study laboratory. Wet fecal samples were collected by the participants (single collection), frozen within 30 min in a sterilized tube and brought to the laboratory immediately. All samples were stored in a −80°C freezer until analysis.

### Animal experiments

All animal studies were performed following the national legislation and was approved by the Institutional Animal Care and Use Committee at the Center for Laboratory Animals, Shanghai Jiao Tong University Affiliated Sixth People’s Hospital (Shanghai, China) and China Agricultural University (Beijing, China).

The pig study was conducted in the Metabolism Laboratory of the National Feed Engineering Technology Research Center (Fengning, Hebei Province, China). Six crossbred growing pigs (Duroc x Landrace x Yorkshire, weighing around 25 kg) were used in this experiment. The pigs were housed individually in stainless steel metabolism cages (1.4 x 0.7 x 0.6 m) equipped with a feeder and a nipple drinker. The crates were located in three environmentally controlled rooms with the temperature maintained at 22-24 ^o^ C. The pigs were allowed a 10-day period to adapt to the metabolism crates and the environment of the room, and were fed commercial corn-soybean meal based diets.

The C57BL/6J mice (male, 6 weeks old) were purchased from Shanghai Laboratory Animal Co Ltd. (Shanghai, China), and the db/db mice inbred on BKS background (male, 8 weeks old) were purchased from Model Animal Research Center of Nanjing University (Nanjing, China). The mousee studies were conducted at the Center for Laboratory Animals, Shanghai Jiao Tong University Affiliated Sixth People’s Hospital (Shanghai, China) after one week of acclimatization. All experimental mice were housed in specific-pathogen-free (SPF) environments under a controlled condition of 12 h light/12 h dark cycle at 20-22 ^o^ C and 45 ± 5 % humidity, with free access to purified rodent diet and ultrapure water. The body weights and the consumption of food and water were measured weekly for the duration of the experiments. The blood glucose levels were measured each week, and OGTT was carried out as described in the results. At the end of each experiment, the retro-orbital blood was collected before sacrifice to measure serum insulin, and GLP-1 concentrations for all of the mice. All samples were stored in a −80°C freezer until analysis.

#### Animal experiment 1: GW4064 treatment in pigs

Six pigs including 3 males and 3 females were used in this experiment. All the pigs were orally administered GW4064 (Hanxiang Corp.) at a dose of 10 mg/kg. The administration was carried out twice with a 12 h interval between doses. Blood samples were collected through a catheter embedded in the precaval vein 15 min, 35 min, 60 min, and 24 h after the second GW4064 administration for BA, blood glucose, and GLP-1 measurements. All samples were stored in a −80°C freezer until analysis.

#### Animal experiment 2: HCA species oral administration in C57BL/6J mice

Twenty C57BL/6J wild type mice were divided into four groups and were orally administrated with the following agents for 28 days: 1) control group: mice (n = 5) were administered with control vehicle, 6 % NaHCO3 (S6014, Sigma-Aldrich); 2) metformin group: mice (n = 5) were administered with metformin (D150959, Sigma-Aldrich) at a daily dose of 200 mg/kg/day; 3) HCA group: mice (n = 5) were administered with HCA (700159P, Sigma-Aldrich) at a daily dose of 100 mg/kg/day; 4) HDCA group: mice (n = 5) were administered with HDCA (H3878, Sigma-Aldrich) at a daily dose of 100 mg/kg/day.

#### Animal experiment 3: HCA species oral administration in HFD+STZ mice

Forty C57BL/6J mice were placed on a high-fat diet (HFD: 60 % kcal from fat; D12492, Research Diets). After 12 weeks of HFD, mice were fasted for 5 h and then injected with a single dose of streptozotocin (STZ; V900890, Sigma-Aldrich) (75 mg/kg i.p.) as a freshly prepared solution in 0.1 mmol/L sodium citrate (S4641, Sigma-Aldrich), pH 5.5. After 72 h post-injection, only STZ-treated mice exhibiting a fasting glucose level ≥11.1 mmol/L were used in the study (n = 32). Thirty-two HFD+STZ mice were divided into four groups and were orally administrated with the following agents for 28 days: 1) control group: mice (n = 8) were administered with control vehicle, 6 % NaHCO_3_; 2) metformin group: mice (n = 8) were administered with metformin at a daily dose of 200 mg/kg/day; 3) HCA group: mice (n = 8) were administered with HCA at a daily dose of 100 mg/kg/day; 4) HDCA group: mice (n = 8) were administered with HDCA at a daily dose of 100 mg/kg/day.

#### Animal experiment 4: HCA oral administration in db/db mice

Twenty-four db/db mice were divided into three groups and were orally administrated with the following agents for 28 days: 1) control group: mice (n = 8) were administered with control vehicle, 6 % NaHCO_3_; 2) metformin group: mice (n = 8) were administered with metformin at a daily dose of 200 mg/kg/day; 3) HCA group: mice (n = 8) were administered with HCA at a daily dose of 100 mg/kg/day.

#### Animal experiment 5: TGR5 antagonist, FXR agonist, and GLP-1 receptor antagonist administration in mice

Forty C57BL/6J mice were divided into eight groups and were administrated with the following agents for 28 days: 1) control group: mice (n = 5) were administered with control vehicle, 6 % NaHCO_3_ (i.g.); 2) HCA group: mice (n = 5) were administered with HCA (100 mg/kg/day, i.g.); 3) 5β-CA group: mice (n = 5) were administered with control vehicle, 6 % NaHCO_3_ (i.g.), and 5β-CA (C7628, Sigma-Aldrich) in 0.5 % Sodium Carboxymethyl Cellulose (CMC-Na; 419338, Sigma-Aldrich) (100 mg/kg/day, i.g.); 4) HCA+5β-CA group: mice (n = 5) were administered with HCA (100 mg/kg/day, i.g.), and 5β-CA in 0.5 % CMC-Na (100 mg/kg/day, i.g.); 5) FEX group: mice (n = 5) were administered with control vehicle, 6 % NaHCO_3_ (i.g.), and FEX (Hanxiang Corp.) in 0.5 % CMC-Na (100 mg/kg/day, i.g.); 6) HCA+FEX group: mice (n = 5) were administered with HCA (100 mg/kg/day, i.g.), and FEX in 0.5 % CMC-Na (100 mg/kg/day, i.g.). 7) Exendin group: mice (n = 5) were administered with control vehicle, 6 % NaHCO_3_ (i.g.), and Exendin in saline (25nmol/kg/day, i.p.); 8) HCA+ Exendin group: mice (n = 5) were administered with HCA (100 mg/kg/day, i.g.), and Exendin (2081, R&D Systems) in saline (25nmol/kg/day, i.p.).

#### Fasting blood glucose measurement and OGTT

Fasting blood glucose measurement and OGTT was carried out in mice after overnight fasting. The glucose levels of tail vain blood samples were analyzed using a glucose analyzer (OneTouch Ultra, Lifescan, Johnson&Johnson, Milpitas, CA). In OGTT, a glucose solution (1.5 g/kg) was orally administered to each mouse, and samples were analyzed for glucose level before (0 min) and at 15min, 30 min, 60 min, and 120 min after the oral glucose load.

#### Serum GLP-1 and insulin measurement

Blood samples were collected and centrifuged at 3,000 x g, 4 ^o^C, for 10 min for serum collection. For GLP-1 analysis, dipeptidyl peptidase IV inhibitor (10 μL/mL; Millipore Corp, Missouri) was added to the blood before serum collection. High sensitivity GLP-1 active chemiluminescent ELISA kit (Millipore Corp, Missouri) and high sensitive mouse insulin immunoassay ELISA kit (ImmunoDiagnostics Limited, Hong Kong) were used for GLP-1 and insulin measurement, respectively.

### Statistical analysis

The BA profile raw data acquired using UPLC-TQ/MS were processed and quantified using TargetLynx software (Waters Corp., Milford, MA). Manual checking and correction were carried out in order to ensure data quality. The HCA species concentration was calculated by combining the concentrations of HCA, HDCA, GHCA, GHDCA, THDCA, and THCA. Non-parametric Mann Whitney U test and Wilcoxon signed-rank test were carried out for comparison of unpaired and paired samples in the human studies. In animal and cell studies, parametric unpaired t-test and paired t test were applied to compare the unpaired and paired samples, respectively. Spearman’s rank correlation coefficients were calculated to examine the association of BAs and typical clinical measurements. ROC (Receiver Operation Curve) analysis was used to test the sensitivity and specificity of total and individual HCA species in group separation. Logistic regression models were constructed to assess the predictive potentials of individual and combined HCA species on future metabolic health. For human studies, the p values were corrected by FDR. For human, animal and cell studies, p<0.05 were considered statistically significant (two tailed). SPSS (V19, IBM, USA), GraphPad Prism (6.0, Graphpad, USA), and MATLAB (2014a, MathWorks, USA) were used for statistical analyses and graphic generation. Analyte levels in tables and figures were presented as mean ± S.E. or mean ± S.D.

Materials and methods on cell studies and quantitative analysis of BAs are provided in supplementary information.

## Acknowledgements

This work was supported by the National Key R&D Program of China (2017YFC0906800), National Natural Science Foundation of China (31501079 31500954 and 81772530), and International Science and Technology Cooperation Program of China (2014DFA31870). We thank the Biobank of Shanghai Sixth People’s Hospital for providing clinical samples for this study.

## Author Contributions

W.J. conceptualized the study and designed the research. X.Z. and T.C. performed the data preprocessing and statistical analysis. W.P.J was the leader of the cohort studies and together with Y.B. provided biospecimens from their studies. W.J., X.Z., T.C., and R.J. drafted the manuscript. W.J., C.R., J.P., X.Z. critically revised the manuscript. A.Z., C.R., J.T., G.X., A.L. and W.Z. provided valuable suggestions in data analysis and interpretation. X.M., Y.B., C.W., H.Y., M.J., A.L., and Y.Y. were responsible for human sample collection and explanation. X.Z., A.H., Y.Z., M.W., M.L., D.L., X.H., F.H., Y.Y., J.L., Q.Z., K.G., S.L., S.W., and Y.L. were responsible for animal sample collection. A.Z., X.Z. and F.H. were responsible for sample preparation and analysis. R.J., M.W. and Q.Z. were responsible for cell studies.

## Author Information

The authors declare that they have no conflicts of interest.

