## Supplementary Information for "Hyocholic acid species and the risk of type 2 diabetes"

**Supplementary material**

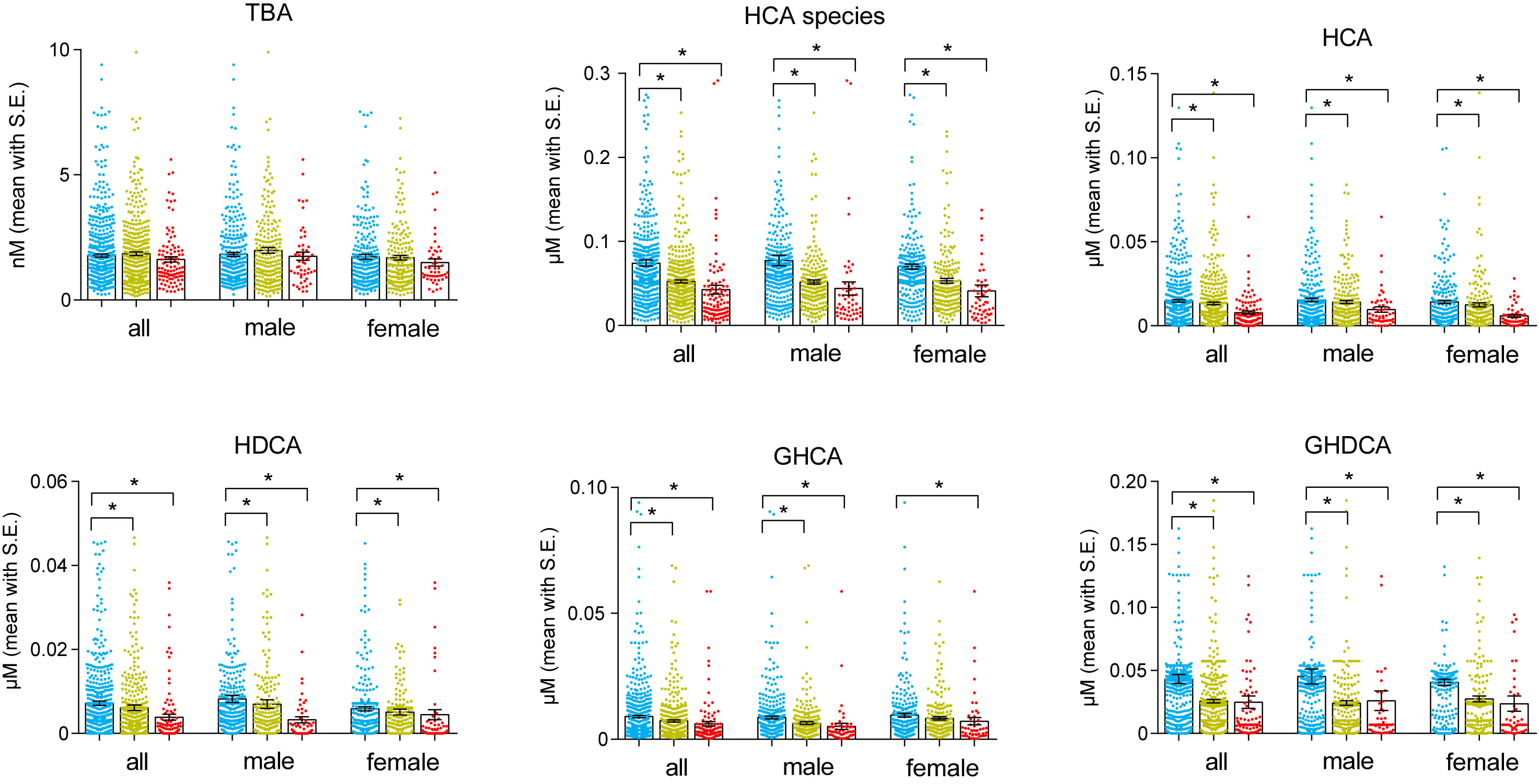

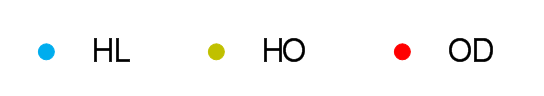

Figure S1 Serum concentrations of total bile acids (TBA) and total and individual HCA species for HL (blue, n=585), HO (yellow, n=419), and OD (red, n=103) groups in the first cross sectional study. Mean with S.E., * FDR corrected Mann-Whitney p<0.05 when compared with HL.

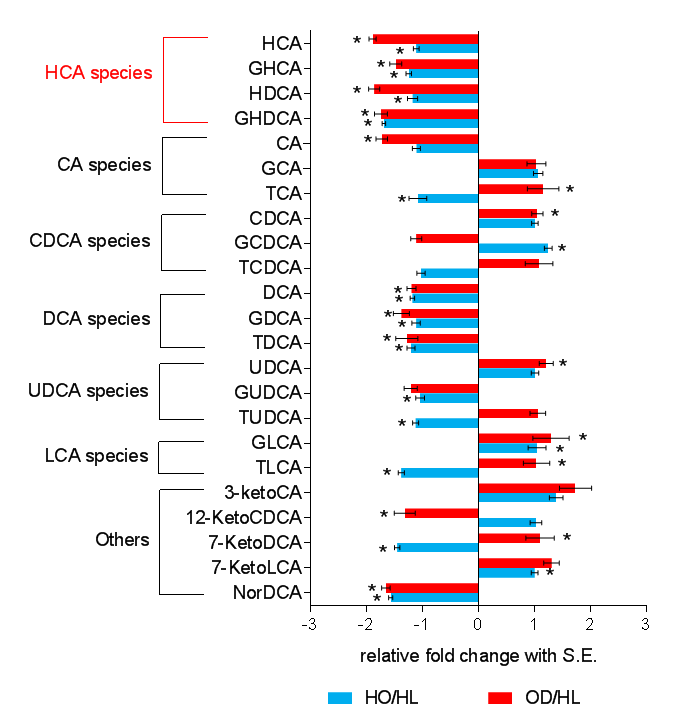

Figure S2 Fold change (mean with S.E.) of 23 BAs in HO and OD groups relative to HL group in the first cross sectional study (all samples). * FDR corrected Mann-Whitney p<0.05 when compared with HL.

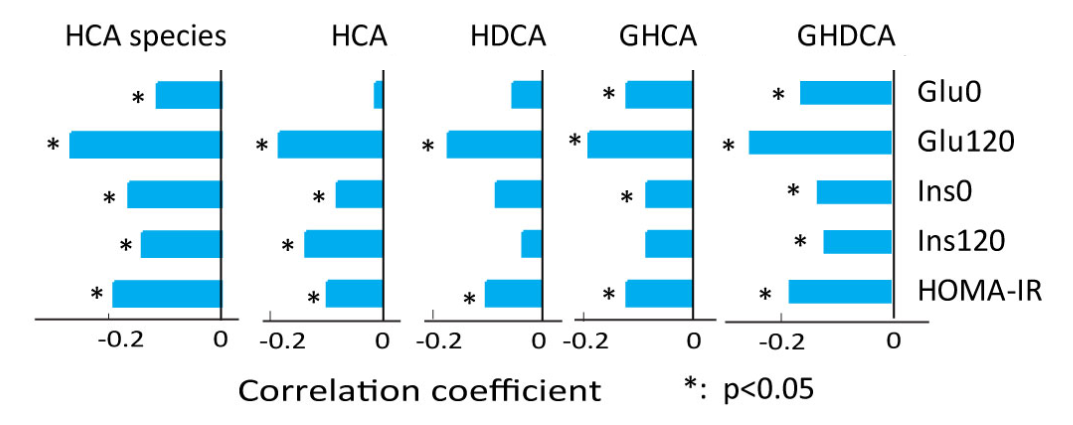

Figure S3 Correlation coefficients of total and individual HCA species with representative glycemic markers in all samples of the first cross sectional study. * p < 0.05.

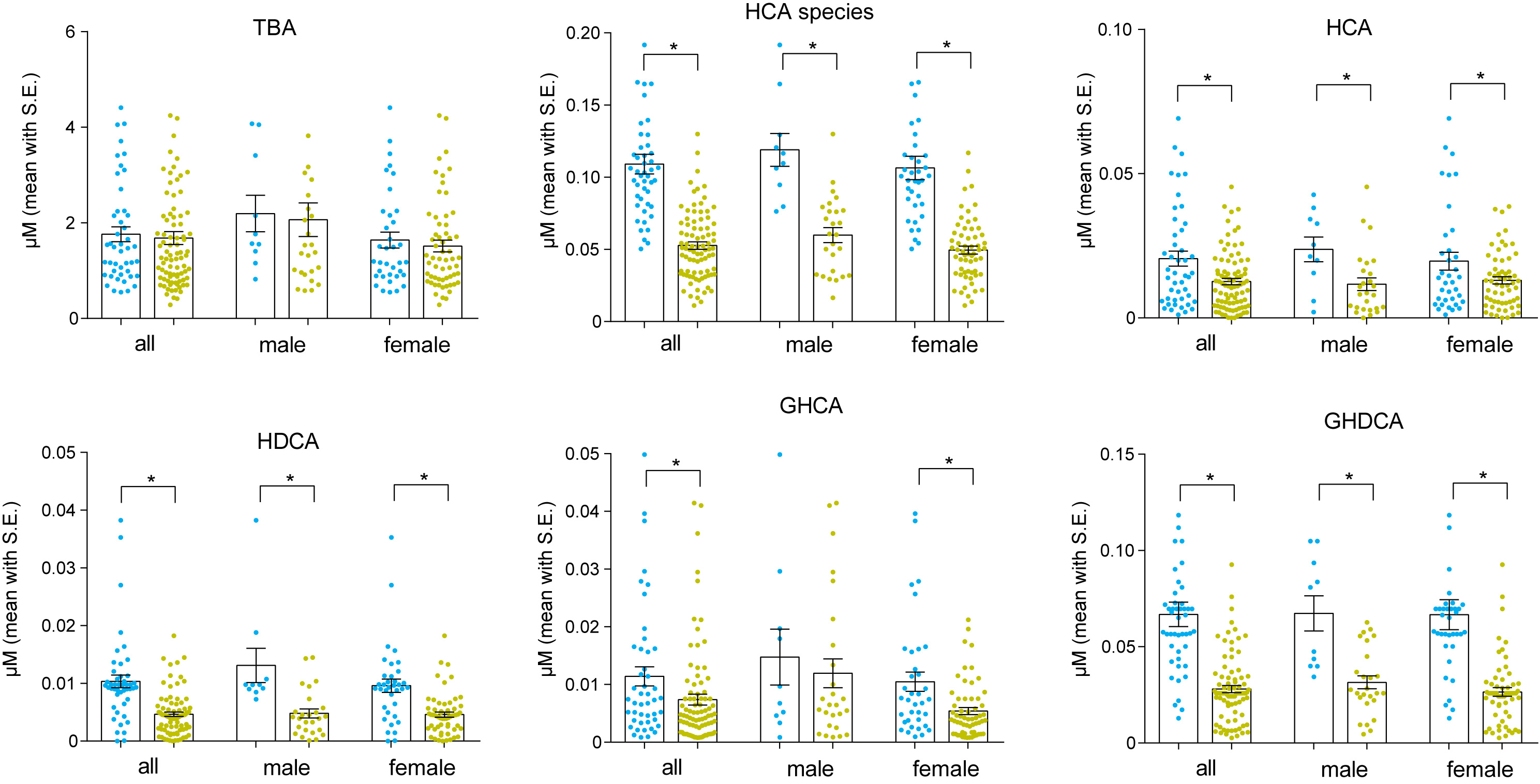

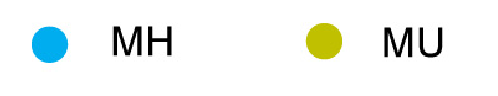

Figure S4 Serum concentrations of total bile acids (TBA) and total and individual HCA species for future metabolically healthy (MH, n=46) and unhealthy (MU, n=86) groups in the 10-year longitudinal study. Mean with S.E., * FDR corrected Mann-Whitney p<0.05 comparing MH and MU.

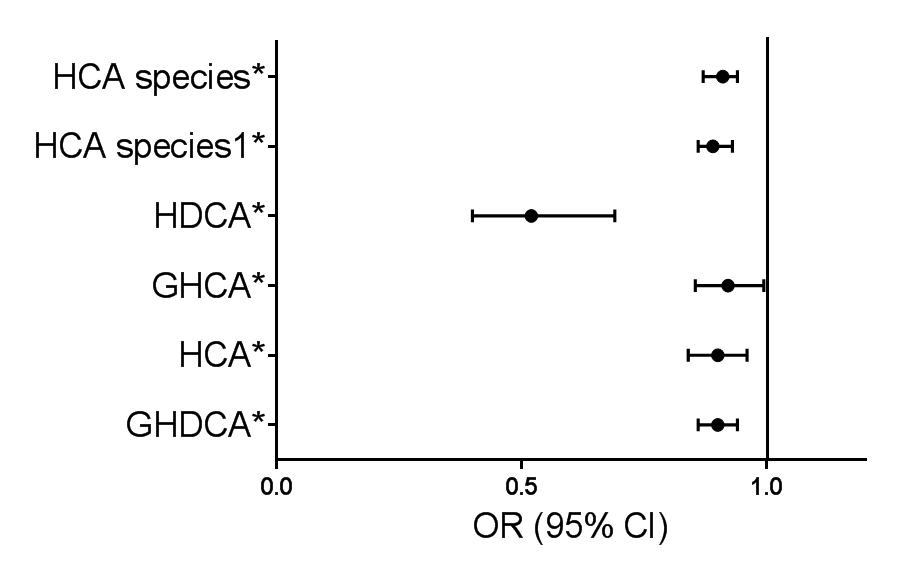

Figure S5	Binary logistic regression (OR with 95% CI) of total and individual HCA species indicates their potential use as risk factor for future metabolic unhealthy outcome in the 10-year longitudinal study. (*p <0.05) HCA species1 represents result of total HCA species after adjustment of age, sex and BMI.

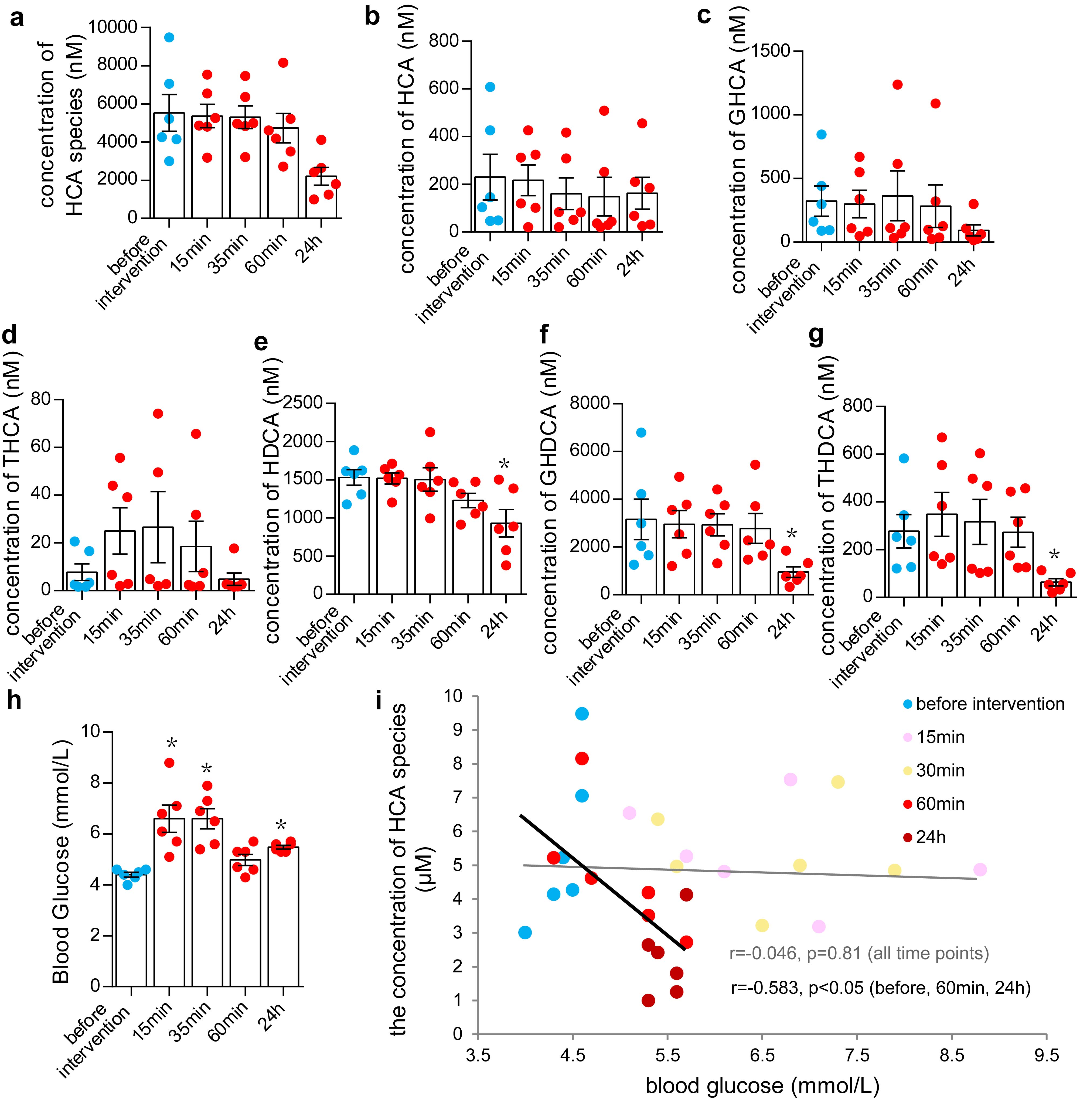
Figure S6 (a-g) Serum concentrations of total and individual HCA species and (h) blood glucose in pigs before and after oral administration of GW4064. Mean with S.E. *p<0.05 when comparing with before intervention using paired t-test.

It should be pointed out that the blood glucose levels were dramatically increased after 15-min and 35-min intervention even higher than those after 24-h intervention (Fig. S12h), while the levels of HCA species rarely changed compared to the baseline level. Such growth of glucose levels might be due to the stressful situation of the pigs shortly after GW4064 administration. To eliminate the stress effect induced glucose increase, the correlation analysis between the glucose levels and the concentration of HCA species was carried out across three time points (before intervention, 60min, and 24h, other than 15min and 35 min, Fig. S12i). The alteration of blood glucose was negatively correlated with HCA species, while such correlation could not be observed across all time points.

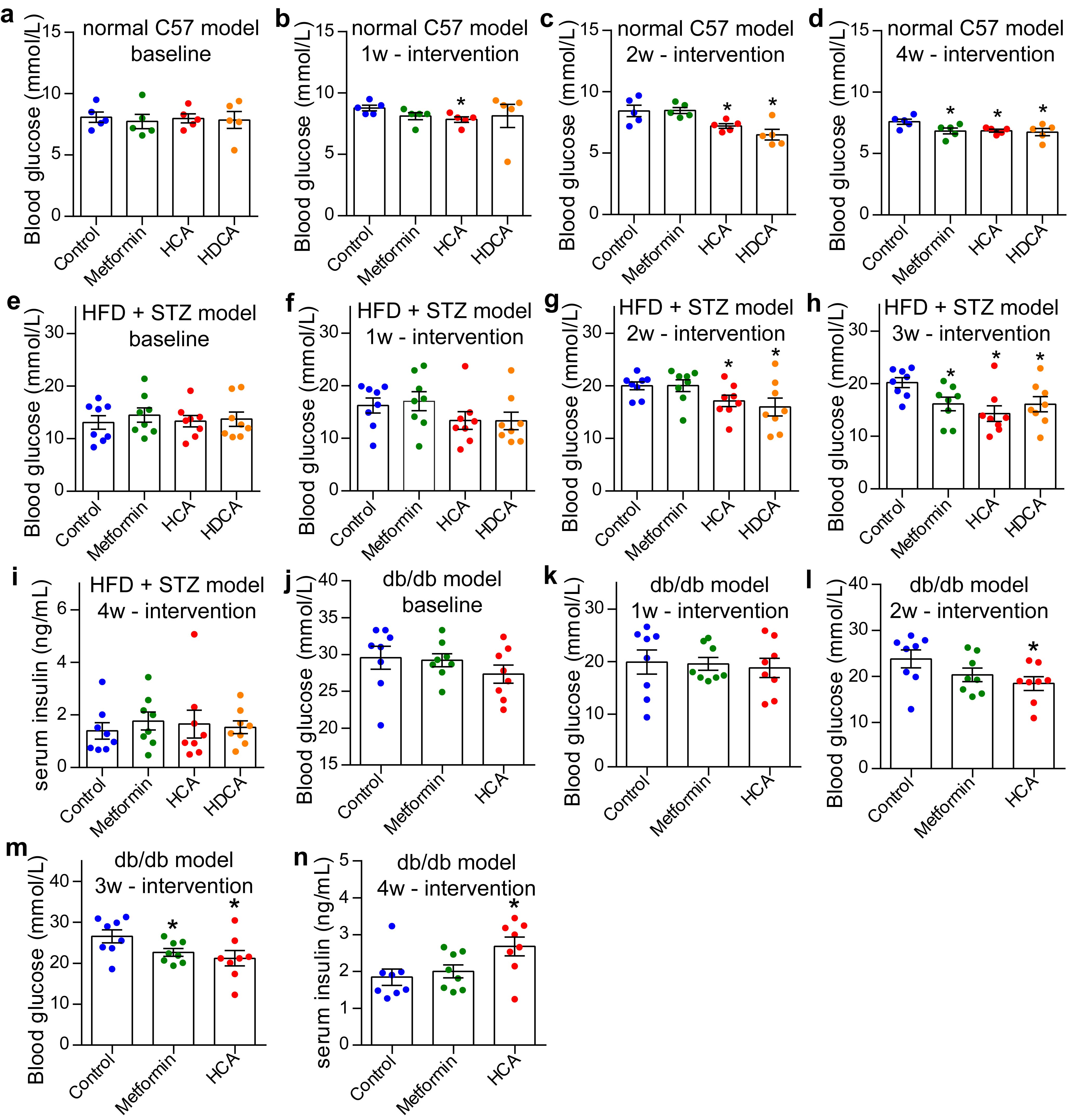

Figure S7 (a-d) Blood glucose levels of the normal C57BL/6J mouse models before intervention and treated with metformin, HCA, HDCA and vehicle control for one to four weeks. (e-h) Blood glucose levels of the HFD+STZ mouse models before intervention and treated with metformin, HCA, HDCA and vehicle control for one to three weeks, and (i) serum insulin levels of HFD+STZ mouse models treated for four weeks. (j-m) Blood glucose levels of the db/db mouse models before intervention and treated with metformin, HCA and vehicle control for one to three weeks, and (n) serum insulin levels of db/db mouse models treated for four weeks. Mean with S.E. * p<0.05 when compared with control group using unpaired t-test.

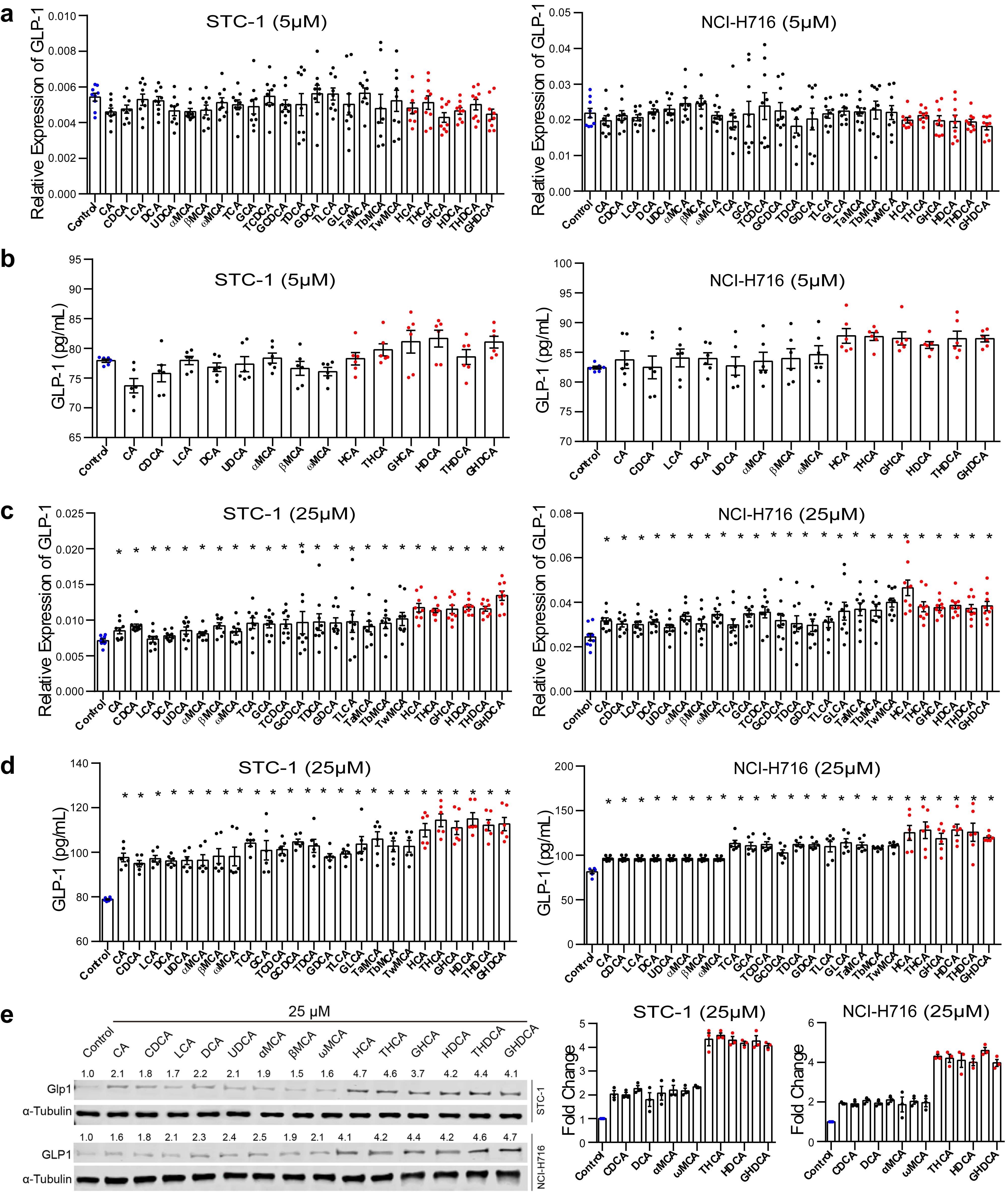

Figure S8 NCI-H716 and STC-1 cells were treated with 6 HCA species and 19 other BAs for 48h with different concentrations (5 and 25 μM). The GLP-1 gene transcription was determined by using Real-time PCR including (a) for 5 μM and (c) for 25 μM. The GLP-1 secretion was determined using ELISA including (b) for 5 μM and (d) for 25 μM. NCI-H716 and STC-1 cells were treated with all 6 HCA species and 8 other representative BAs, each at 25 µM for 48 h, and the intracellular GLP-1 protein expression was measured using western-blot (e). Representative images are shown, and data were obtained from 3 independent experiments. Mean with S.E. * p<0.05 when compared with control using unpaired t-test.

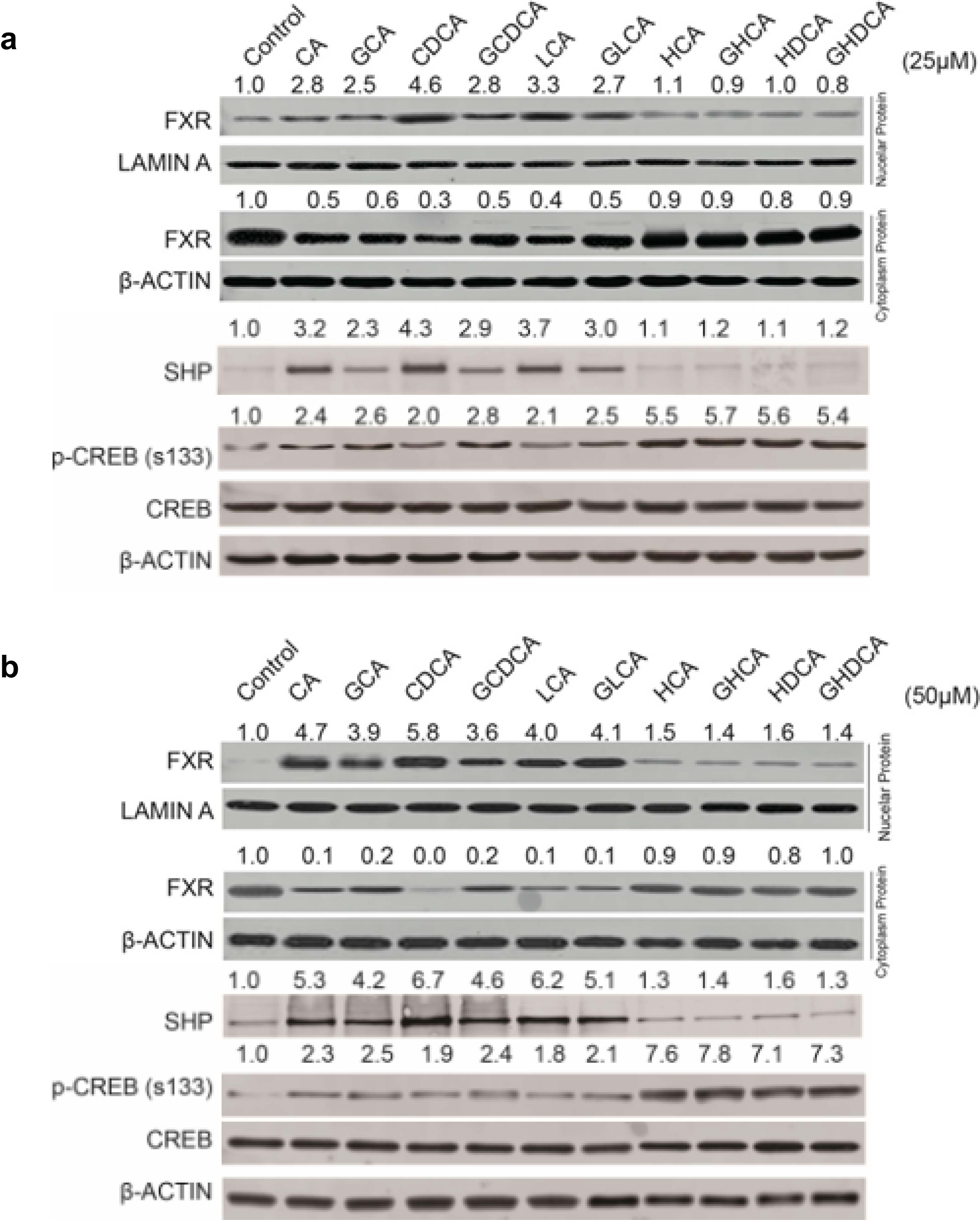

Figure S9 FXR protein concentration in nuclear and cytosolic fractions, FXR downstream protein SHP, p-CREB, total CREB of NCI-H716 cells treated with 25 μM (a) and 50 μM (b) of 4 representative HCA species and 6 other representative BAs for 24 hours.

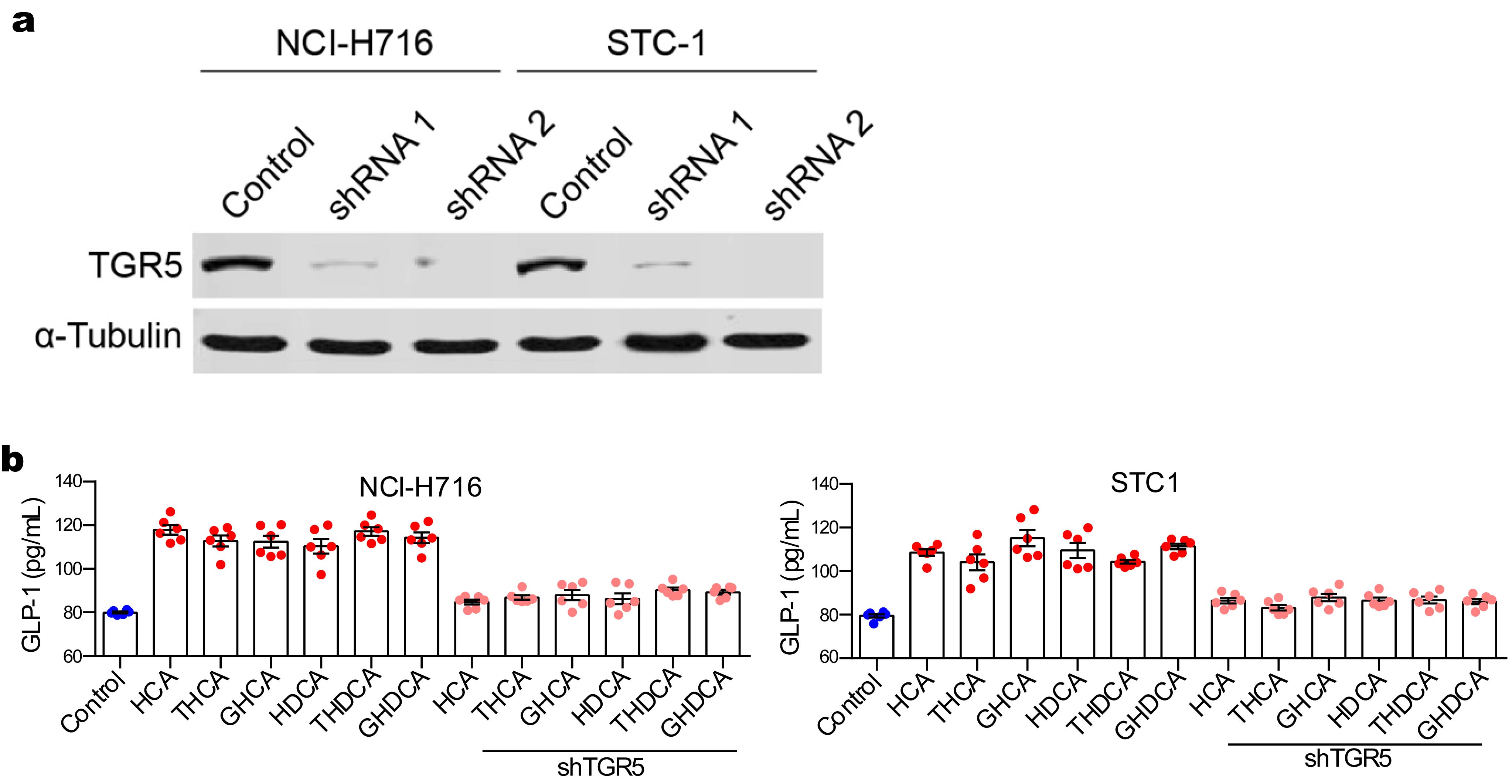

Figure S10 (a) TGR5 expression in NCI-H716 and STC-1 was knocked down by using two specific shRNAs, and the expression of TGR5 was determined using western-blot. (b) NCI-H716 and STC-1 as well as their TGR5 knockdown cells were treated with 6 HCA species for 24h, and the GLP-1 secretion was determined using ELISA. Mean with S.E.

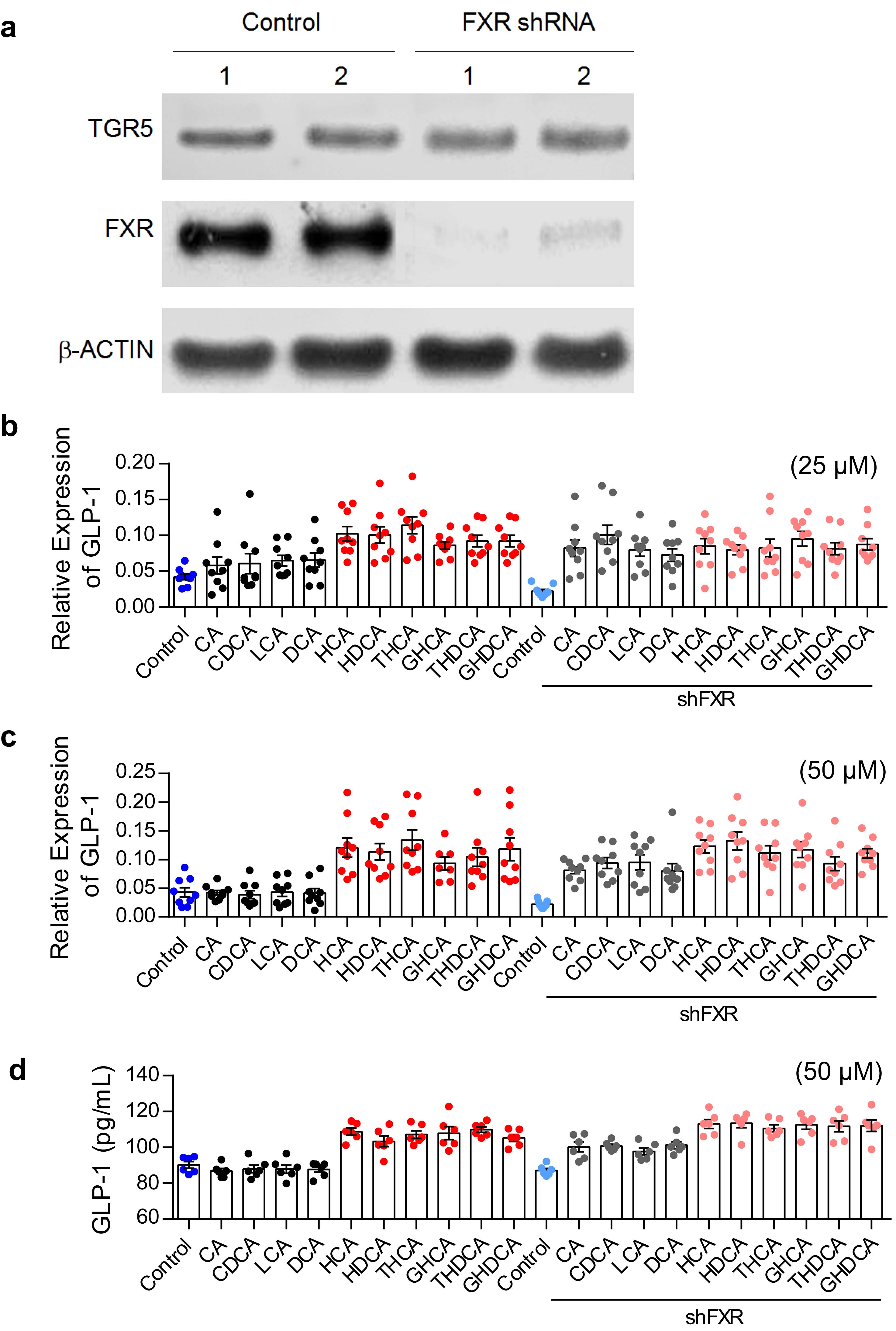

Figure S11 (a) FXR expression in NCI-H716 cells was knocked down by using a specific shRNA, and the expression of TGR5 and FXR was determined using western-blot. NCI-H716 cells as well as their FXR knockdown cells were treated with 6 HCA species and 4 other representative BAs at the concentration of 25 μM and 50 μM for 24h. The GLP-1 transcription was measured using Real-time PCR at 25 μM (b) and 50 μM (c). The GLP-1 secretion was measured using ELISA at 50 μM (d). Mean with S.E.

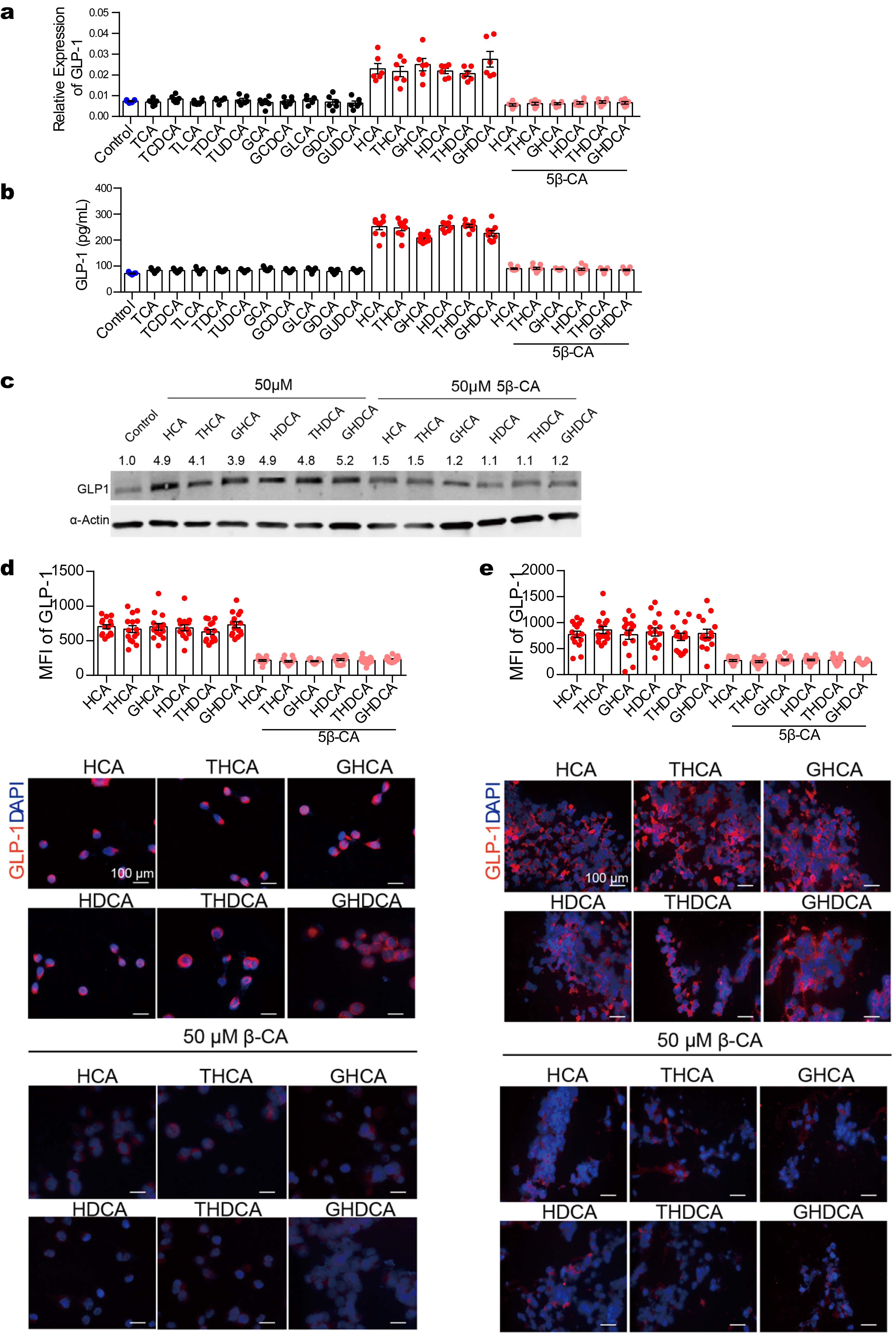

Figure S12 (a, b) GLP-1 gene expression (a, RT-qPCR) and secretion (b, ELISA) in NCI-H716 cells treated with 50 μM HCA species or other BAs for 48 hours, with or without 50 μM 5β-CA. (c-e) Quantification of intracellular GLP-1 protein expression using western blot (c), IF staining in 2D (d, x200) and 3D (e, x200) cultured NCI-H716 cells treated 50 μM HCA species for 48 hours (2D culture) or 7 days (3D culture), with or without 50 μM 5β-CA. Representative images are shown, data obtained from 3 independent experiments. The IF staining images were analyzed using Image J for Mean Florescence Intensity (MFI). Mean with S.E.

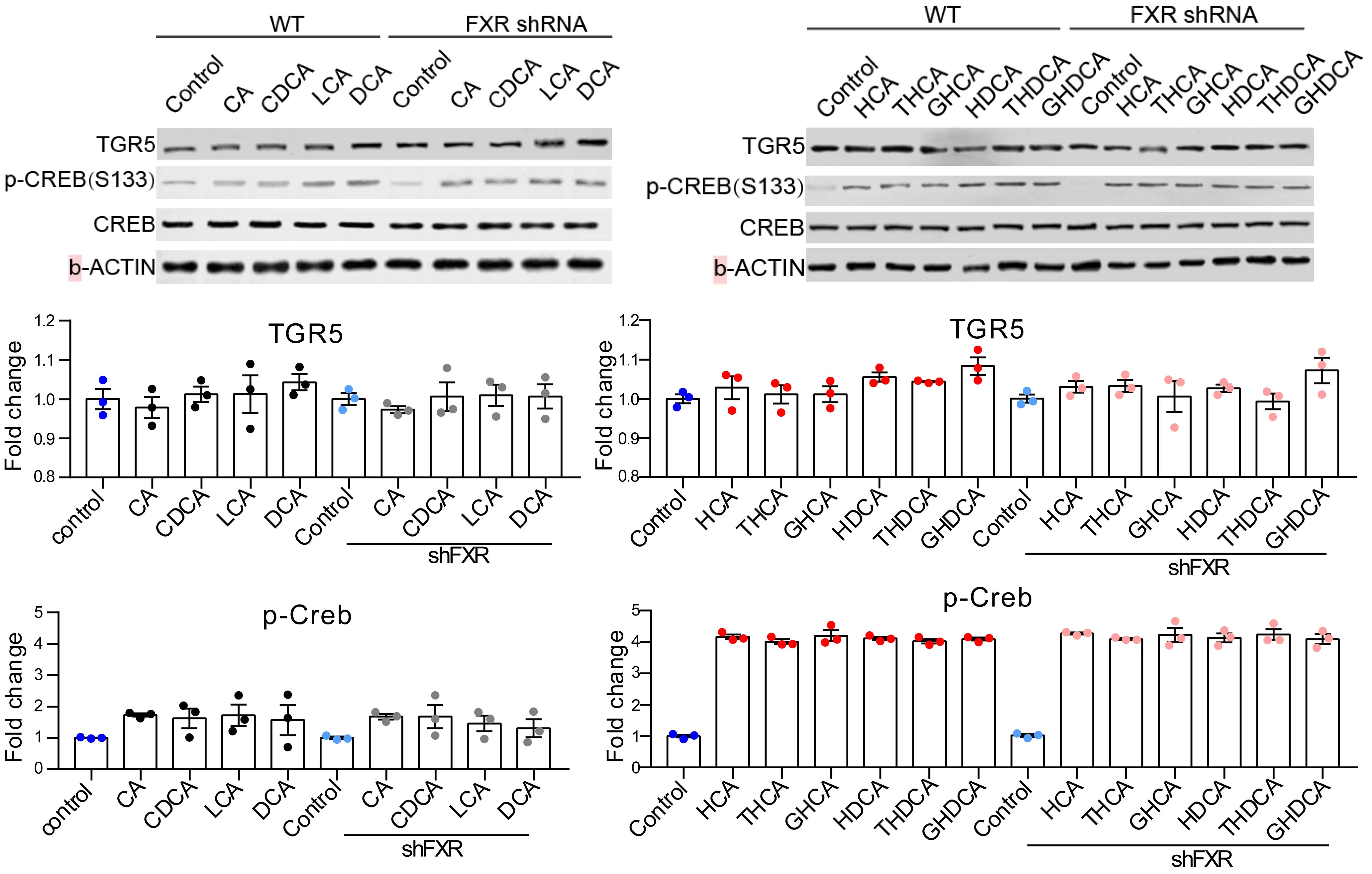

Figure S13 NCI-H716 cells and their FXR knockdown cells were treated with 6 HCA species and 4 representative BAs for 24h, and TGR5, p-CREB and total CREB were determined using western blot. Representative images are shown, and data were obtained from 3 independent experiments. Mean with S.E.

Table S1. Metabolic markers of all healthy lean (HL), healthy overweight/obese (HO) and overweight/obese with type 2 diabetes (OD) subjects in the first cross sectional study

|  | Healthy lean (HL) | Healthy overweight/obese (HO) | Overweight/obese with type 2 diabetes (OD) |
| --- | --- | --- | --- |
| n(M/F) | 585 (329/256) | 419 (229/190) | 103 (52/51) |
| Age (year) | 37.1±11.7 | 39.8±11.8* | 53.4±10.0* |
| BMI (kg/m2) | 21.3±1.7 | 28.3±3.4* | 29.0±3.5* |
| Glu0 (mmol/L) | 5.1±0.5 | 5.2±0.6 | 8.2±2.5* |
| Glu120 (mmol/L) | 5.3±1.1 | 6.1±1.4* | 14.4±4.0* |
| Ins0 (mU/L) | 5.9±2.7 | 11.3±10.9* | 16.4±19.4* |
| Ins120 (mU/L) | 30.6±19.3 | 61.8±59.2* | 76.4±57.4* |
| TC (mmol/L) | 4.8±0.9 | 4.7±0.9 | 6.2±1.7* |
| TG (mmol/L) | 1.2±0.6 | 1.4±0.8* | 2.5±2.4* |
| HDL (mmol/L) | 1.5±0.3 | 1.4±0.3* | 1.2±0.2* |
| LDL (mmol/L) | 2.5±0.5 | 2.7±0.5* | 3.2±0.8* |
| SP (mmHg) | 112.6±11.1 | 116.4±13.1* | 137.8±20.2* |
| DP (mmHg) | 72.0±7.2 | 75.0±8.6* | 83.5±13.4* |
| HeartRate (bpm) | 75.7±7.7 | 75.7±6.6* | 78.4±6.1* |
| ALT (U/L) | 24.9±14.2 | 30.1±17.8* | 28.9±19.6 |
| AST (U/L) | 20.8±5.7 | 22.1±7.3* | 23.4±11.5 |
| HOMA-IR | 1.3±0.6 | 2.6±3.0* | 6.0±7.8* |

Values were presented as number or mean ± S.D.

* Mann Whitney p<0.05 when compared with HL. Chi-Square was used to compare sex ratio between groups.

BMI = body mass index; SP = systolic [blood](http://cn.bing.com/dict/clientsearch?mkt=zh-CN&setLang=zh&form=BDVEHC&q=收缩压) [pressure](http://cn.bing.com/dict/clientsearch?mkt=zh-CN&setLang=zh&form=BDVEHC&q=收缩压); DP = diastolic [blood](http://cn.bing.com/dict/clientsearch?mkt=zh-CN&setLang=zh&form=BDVEHC&q=收缩压) [pressure](http://cn.bing.com/dict/clientsearch?mkt=zh-CN&setLang=zh&form=BDVEHC&q=收缩压); Glu0 = fasting plasma glucose; Glu120 = 2 h plasma glucose; Ins0 = fasting insulin; Ins120 = 2 h insulin; TG = total triglycerides; TC = total cholesterol; HDL = high density lipoprotein; LDL = low density lipoprotein; ALT = alanine aminotransferase; AST = aspartate aminotransferase; HOMA-IR = Glu0×Ins0/22.5.

Table S2. Metabolic markers of matched healthy lean (HL), healthy overweight/obese (HO) and overweight/obese with type 2 diabetes (DM) subjects in the first cross sectional study

|  | Healthy lean (HL) | Healthy overweight/obese (HO) | Overweight/obese with type 2 diabetes (OD) |
| --- | --- | --- | --- |
| n(M/F) | 103 (52/51) | 103 (52/51) | 103 (52/51) |
| Age (year) | 53.8±5.6 | 52.9±10.1 | 53.4±10.0 |
| BMI (kg/m2) | 21.6±1.8 | 28.7±4.4* | 29.0±3.5* |
| Glu0 (mmol/L) | 5.3±0.6 | 5.3±0.9 | 8.2±2.5* |
| Glu120 (mmol/L) | 5.6±1.2 | 6.1±1.4 | 14.4±4.0* |
| Ins0 (mU/L) | 5.2±2.5 | 11.6±11.9* | 16.4±19.4* |
| Ins120 (mU/L) | 28.1±18.5 | 52.9±55.5* | 76.4±57.4* |
| TC (mmol/L) | 5.2±1.1 | 4.9±1.1* | 6.2±1.7* |
| TG (mmol/L) | 1.3±0.6 | 1.3±0.6 | 2.5±2.4* |
| HDL (mmol/L) | 1.7±0.3 | 1.4±0.3* | 1.2±0.2* |
| LDL (mmol/L) | 2.6±0.5 | 2.8±0.4* | 3.2±0.8* |
| SP (mmHg) | 116.0±10.4 | 119.5±15.1 | 137.8±20.2* |
| DP (mmHg) | 71.8±6.6 | 75.3±10.6* | 83.5±13.4* |
| HeartRate (bpm) | 73.8±7.1 | 75.1±6.7 | 78.4±6.1* |
| ALT (U/L) | 23.5±12.5 | 25.4±12.4 | 29.0±19.6 |
| AST (U/L) | 21.1±5.3 | 21.8±4.6 | 23.4±11.5 |
| HOMA-IR | 1.2±0.5 | 2.7±3.3* | 6.0±7.8* |

Values were presented as number or mean ± S.D.

* Mann Whitney p<0.05 when compared with HL. Chi-Square was used to compare sex ratio between groups.

BMI = body mass index; SP = [systolic](http://cn.bing.com/dict/clientsearch?mkt=zh-CN&setLang=zh&form=BDVEHC&q=收缩压) [blood](http://cn.bing.com/dict/clientsearch?mkt=zh-CN&setLang=zh&form=BDVEHC&q=收缩压) [pressure](http://cn.bing.com/dict/clientsearch?mkt=zh-CN&setLang=zh&form=BDVEHC&q=收缩压); DP = diastolic [blood](http://cn.bing.com/dict/clientsearch?mkt=zh-CN&setLang=zh&form=BDVEHC&q=收缩压) [pressure](http://cn.bing.com/dict/clientsearch?mkt=zh-CN&setLang=zh&form=BDVEHC&q=收缩压); Glu0 = fasting plasma glucose; Glu120 = 2 h plasma glucose; Ins0 = fasting insulin; Ins120 = 2 h insulin; TG = total triglycerides; TC = total cholesterol; HDL = high density lipoprotein; LDL = low density lipoprotein; ALT = alanine aminotransferase; AST = aspartate aminotransferase; HOMA-IR = Glu0×Ins0/22.5.

Table S3. Metabolic markers of healthy control, pre-diabetic and diabetic subjects in the second cross sectional study (serum samples)

|  | Healthy control (C) | Pre-diabetic (Pre) | Diabetic (DM) |
| --- | --- | --- | --- |
| n(M/F) | 32 (12/20) | 34 (12/22) | 40 (20/20) |
| Age (year) | 64.0±11.5 | 64.3±10.4 | 62.5±9.4 |
| BMI (kg/m2) | 23.0±4.2 | 24.6±3.3 | 25.9±3.3* |
| HbA1c (%) | 4.9±0.6 | 5.8±0.5* | 6.5±1.9* |
| Waist (cm) | 82.8±11.4 | 90.1±11.0* | 91.1±10.9* |
| SP (mmHg) | 125.0±8.9 | 132.4±13.7 | 128.3±10.8 |
| DP (mmHg) | 85.9±28.7 | 80.1±9.3 | 79.7±9.3 |
| Glu0 (mmol/L) | 5.0±0.4 | 6.2±0.6* | 8.2±4.4* |
| Glu120 (mmol/L) | 6.7±0.7 | 8.5±1.2* | 12.9±1.8* |
| Ins0 (mU/L) | 8.8±4.4 | 8.7±5.2 | 12.5±6.7 |
| Ins120 (mU/L) | 61.8±28.9 | 99.5±46.4 | 94.7±79.0 |
| HOMA-IR | 2.1±1.0 | 2.1±1.3 | 3.8±2.2 |

Values were presented as number or mean ± S.D..

* Mann Whitney p<0.05 when compared with healthy control. Chi-Square was used to compare sex ratio between groups.

BMI = body mass index; SP = [systolic](http://cn.bing.com/dict/clientsearch?mkt=zh-CN&setLang=zh&form=BDVEHC&q=收缩压) [blood](http://cn.bing.com/dict/clientsearch?mkt=zh-CN&setLang=zh&form=BDVEHC&q=收缩压) [pressure](http://cn.bing.com/dict/clientsearch?mkt=zh-CN&setLang=zh&form=BDVEHC&q=收缩压); DP = diastolic [blood](http://cn.bing.com/dict/clientsearch?mkt=zh-CN&setLang=zh&form=BDVEHC&q=收缩压) [pressure](http://cn.bing.com/dict/clientsearch?mkt=zh-CN&setLang=zh&form=BDVEHC&q=收缩压); Glu0 = fasting plasma glucose; Glu120 = 2 h plasma glucose; Ins0 = fasting insulin; Ins120 = 2 h insulin; HOMA-IR = Glu0×Ins0/22.5.

Table S4. Metabolic markers of healthy control, pre-diabetic and diabetic subjects in the second cross sectional study (fecal samples)

|  | Healthy control (C) | Pre-diabetic (Pre) | Diabetic (DM) |
| --- | --- | --- | --- |
| n(M/F) | 26 (9/17) | 30 (10/20) | 35 (16/19) |
| Age (year) | 58.7±12.6 | 61.6±13.1 | 63±8.5 |
| BMI (kg/m2) | 23.1±4.5 | 24.65±3.19 | 25.72±3.46* |
| HbA1c (%) | 4.9±0.6 | 5.77±0.47* | 6.49±1.99* |
| Waist (cm) | 82.5±11.8 | 90.07±10.75 | 90.47±11.07* |
| SP (mmHg) | 123.6±7.9 | 131.63±15.15 | 127.44±11.83 |
| DP (mmHg) | 89.9±31.4 | 77.88±9.92 | 77.22±8.29 |
| Glu0 (mmol/L) | 5.0±0.4 | 6.2±0.6* | 8.1±4.5* |
| Glu120 (mmol/L) | 6.7±0.6 | 8.4±1.2* | 13.2±1. 5* |
| Ins0 (mU/L) | 9.7±4.5 | 10.0±4.7 | 10.9±5.4 |
| Ins120 (mU/L) | 68.9±28.0 | 102.3±48.5 | 91.3±60.9 |
| HOMA-IR | 2.3±1.0 | 2.4±1.2 | 3.3±2.0 |

Values were presented as number or mean ± S.D..

* Mann Whitney p<0.05 when compared with healthy control. Chi-Square was used to compare sex ratio between groups.

BMI = body mass index; SP = [systolic](http://cn.bing.com/dict/clientsearch?mkt=zh-CN&setLang=zh&form=BDVEHC&q=收缩压) [blood](http://cn.bing.com/dict/clientsearch?mkt=zh-CN&setLang=zh&form=BDVEHC&q=收缩压) [pressure](http://cn.bing.com/dict/clientsearch?mkt=zh-CN&setLang=zh&form=BDVEHC&q=收缩压); DP = diastolic [blood](http://cn.bing.com/dict/clientsearch?mkt=zh-CN&setLang=zh&form=BDVEHC&q=收缩压) [pressure](http://cn.bing.com/dict/clientsearch?mkt=zh-CN&setLang=zh&form=BDVEHC&q=收缩压); Glu0 = fasting plasma glucose; Glu120 = 2 h plasma glucose; Ins0 = fasting insulin; Ins120 = 2 h insulin; HOMA-IR = Glu0×Ins0/22.5.

Table S5. Serum bile acid concentration (μM) of healthy control (C), pre-diabetic (Pre) and diabetic (DM) subjects in the second cross sectional study

|  | Healthy control (C) | Pre-diabetes (Pre) | Diabetes (DM) |
| --- | --- | --- | --- |
| HCA | 1.59E-02±2.64E-03 | 1.13E-02±1.31E-03 | 9.71E-03±1.48E-03* |
| GHCA | 1.09E-02±1.34E-03 | 6.43E-03±7.20E-04* | 6.98E-03±6.60E-04* |
| HDCA | 7.47E-03±7.80E-04 | 5.33E-03±2.80E-04 | 6.10E-03±5.00E-04 |
| GHDCA | 4.07E-02±5.60E-03 | 2.90E-02±4.16E-03 | 2.82E-02±3.75E-03 |
| 12-ketoCDCA | 4.21E-02±9.10E-03 | 3.06E-02±1.13E-02 | 4.93E-02±1.53E-02 |
| 3-ketoCA | 6.11E-03±2.08E-03 | 6.06E-03±1.11E-03 | 3.63E-03±6.80E-04 |
| 7-ketoDCA | 6.32E-03±1.58E-03 | 5.72E-03±1.03E-03 | 4.73E-03±4.60E-04 |
| 7-ketoLCA | 1.19E-02±2.01E-03 | 1.20E-02±2.81E-03 | 1.66E-02±5.19E-03 |
| CA | 8.37E-02±1.94E-02 | 1.42E-01±3.16E-02 | 1.20E-01±2.68E-02 |
| CDCA | 3.12E-01±6.19E-02 | 3.29E-01±5.35E-02 | 2.49E-01±3.65E-02 |
| DCA | 1.48E-01±3.19E-02 | 1.15E-01±1.88E-02 | 1.51E-01±1.74E-02 |
| GCA | 1.84E-01±2.78E-02 | 1.24E-01±1.56E-02 | 1.57E-01±2.21E-02 |
| GDCA | 1.40E-01±2.63E-02 | 7.42E-02±1.28E-02 | 1.42E-01±2.44E-02 |
| GLCA | 5.99E-03±2.04E-03 | 4.04E-03±1.27E-03 | 7.18E-03±2.08E-03 |
| GUDCA | 8.58E-02±1.62E-02 | 5.57E-02±1.40E-02 | 4.95E-02±1.03E-02* |
| NorDCA | 2.58E-03±3.30E-04 | 2.51E-03±2.80E-04 | 2.33E-03±1.70E-04 |
| TLCA | 2.44E-03±2.30E-04 | 2.20E-03±2.20E-04 | 4.49E-03±9.00E-04 |
| UDCA | 9.83E-02±1.86E-02 | 1.14E-01±2.25E-02 | 8.90E-02±1.85E-02 |
| GCDCA | 5.34E-01±6.20E-02 | 5.38E-01±4.67E-02 | 4.18E-01±6.53E-02 |
| TCA | 2.44E-02±4.70E-03 | 1.39E-02±2.58E-03 | 2.39E-02±4.85E-03 |
| TCDCA | 4.65E-02±6.44E-03 | 3.13E-02±6.34E-03* | 5.33E-02±1.12E-02 |
| TDCA | 2.14E-02±3.79E-03 | 1.11E-02±1.89E-03* | 2.33E-02±4.06E-03 |
| TUDCA | 4.43E-03±8.50E-04 | 4.82E-03±2.84E-03* | 4.84E-03±2.06E-03 |

Values were presented as mean ± S.E.

* p<0.05 when compared with healthy control, Mann-Whitney test, after FDR correction.

Table S6. Fecal bile acid concentration (nmol/g) of healthy control (C), pre-diabetic (Pre) and diabetic (DM) subjects in the second cross sectional study

|  | Healthy control (C) | Pre-diabetes (Pre) | Diabetes (DM) |
| --- | --- | --- | --- |
| HCA | 8.61±1.08 | 3.68±0.61* | 3.53±0.62* |
| HDCA | 7.80±0.68 | 3.77±0.51* | 3.34±0.47* |
| norDCA | 0.39±0.25 | 0.33±0.10 | 0.37±0.11 |
| 7-ketoLCA | 444.13±172.60 | 526.07±118.73 | 239.72±86.15 |
| UDCA | 404.49±126.26 | 772.15±176.65 | 443.04±139.37 |
| CDCA | 271.01±89.17 | 374.31±92.73 | 330.12±131.29 |
| DCA | 609.22±111.14 | 870.12±275.75 | 1252.75±351.54 |
| 7-ketoDCA | 142.36±52.85 | 241.12±68.7 | 100.55±36.58 |
| 12-ketoCDCA | 1.30±0.62 | 0.61±0.10 | 0.98±0.24 |
| 3-ketoCA | 62.61±25.12 | 88.67±25.90 | 35.26±14.91 |
| CA | 1022.92±284.20 | 827.46±206.73 | 842.32±253.29 |
| GUDCA | 3.82±1.45 | 6.75±1.90 | 2.07±0.52 |
| GCDCA | 15.23±3.06 | 40.45±11.38 | 29.99±12.44 |
| GDCA | 31.52±22.37 | 21.76±8.90 | 14.82±3.86 |
| GCA | 7.50±1.50 | 15.47±3.42 | 14.28±4.80 |
| TCA | 14.14±6.20 | 22.09±10.03 | 15.09±6.71 |
| LCA | 1845.04±233.77 | 1785.03±259.04 | 2173.80±231.64 |
| TLCA | 17.25±15.22 | 2.74±0.68 | 1.81±0.19 |
| GLCA | 1.29±0.46 | 1.41±0.61 | 1.49±0.42 |

Values were presented as mean ± S.E..

* p<0.05 when compared with healthy control, Mann-Whitney test, after FDR correction.

Table S7. Baseline metabolic markers of all future metabolically healthy and unhealthy groups in the 10-year longitudinal study

|  | future metabolically healthy (MH) | future metabolically unhealthy (MU) |
| --- | --- | --- |
| n(M/F) | 46 (10/36) | 86 (26/60) |
| Age (year) | 32.8±9.6 | 39.7±11.7* |
| BMI (kg/m2) | 23.0±2.8 | 24.6±3.5* |
| Waist (cm) | 74.1±9.0 | 79.7±9.9* |
| Glu0 (mmol/L) | 4.7±0.4 | 4.7±0.4 |
| Glu120 (mmol/L) | 4.6±0.9 | 5.1±1.1 |
| Ins0 (mU/L) | 6.3±2.7 | 6.6±3.2 |
| Ins120 (mU/L) | 31.3±25.0 | 34.8±22.5 |
| HDL (mmol/L) | 1.4±0.2 | 1.4±0.2 |
| LDL (mmol/L) | 2.5±0.5 | 2.7±0.5 |
| SP (mmHg) | 109.3±11.7 | 112.6±11.9 |
| DP (mmHg) | 71.4±6.5 | 73.2±6.2 |
| HOMA-IR | 1.3±0.6 | 1.4±0.7 |

Values were presented as number or mean ± S.D..

* Mann Whitney p<0.05 when comparing the two groups, after FDR correction. Chi-Square was used to compare sex ratio between groups.

BMI = body mass index; SP = [systolic](http://cn.bing.com/dict/clientsearch?mkt=zh-CN&setLang=zh&form=BDVEHC&q=收缩压) [blood](http://cn.bing.com/dict/clientsearch?mkt=zh-CN&setLang=zh&form=BDVEHC&q=收缩压) [pressure](http://cn.bing.com/dict/clientsearch?mkt=zh-CN&setLang=zh&form=BDVEHC&q=收缩压); DP = diastolic [blood](http://cn.bing.com/dict/clientsearch?mkt=zh-CN&setLang=zh&form=BDVEHC&q=收缩压) [pressure](http://cn.bing.com/dict/clientsearch?mkt=zh-CN&setLang=zh&form=BDVEHC&q=收缩压); Glu0 = fasting plasma glucose; Glu120 = 2 h plasma glucose; Ins0 = fasting insulin; Ins120 = 2 h insulin; HDL = high-density lipoprotein; LDL = low-density lipoprotein; HOMA-IR = Glu0×Ins0/22.5.

Table S8. Baseline metabolic markers of matched future metabolically healthy and unhealthy groups in the 10-year longitudinal study

|  | future metabolically healthy (MH) | future metabolically unhealthy (MU) |
| --- | --- | --- |
| n(M/F) | 46 (10/36) | 46 (10/36) |
| Age (year) | 32.8±9.6 | 33.2±6.2 |
| BMI (kg/m2) | 23.0±2.8 | 22.3±2.1 |
| Waist (cm) | 74.1±9.0 | 74.1±7.3 |
| Glu0 (mmol/L) | 4.7±0.4 | 4.7±0.4 |
| Glu120 (mmol/L) | 4.6±0.9 | 4.9±1.1 |
| Ins0 (mU/L) | 6.3±2.7 | 6.3±2.9 |
| Ins120 (mU/L) | 31.3±25.0 | 31.8±21.3 |
| HDL (mmol/L) | 1.4±0.2 | 1.4±0.2 |
| LDL (mmol/L) | 2.5±0.5 | 2.6±0.5 |
| SP (mmHg) | 109.3±11.7 | 109.4±11.0 |
| DP (mmHg) | 71.4±6.5 | 72.1±7.0 |
| HOMA-IR | 1.3±0.6 | 1.3±0.7 |

Values were presented as number or mean ± S.D..

* Mann Whitney p<0.05 when comparing the two groups, after FDR correction. Chi-Square was used to compare sex ratio between groups.

BMI = body mass index; SP = [systolic](http://cn.bing.com/dict/clientsearch?mkt=zh-CN&setLang=zh&form=BDVEHC&q=收缩压) [blood](http://cn.bing.com/dict/clientsearch?mkt=zh-CN&setLang=zh&form=BDVEHC&q=收缩压) [pressure](http://cn.bing.com/dict/clientsearch?mkt=zh-CN&setLang=zh&form=BDVEHC&q=收缩压); DP = diastolic [blood](http://cn.bing.com/dict/clientsearch?mkt=zh-CN&setLang=zh&form=BDVEHC&q=收缩压) [pressure](http://cn.bing.com/dict/clientsearch?mkt=zh-CN&setLang=zh&form=BDVEHC&q=收缩压); Glu0 = fasting plasma glucose; Glu120 = 2 h plasma glucose; Ins0 = fasting insulin; Ins120 = 2 h insulin; HDL = high-density lipoprotein; LDL = low-density lipoprotein; HOMA-IR = Glu0×Ins0/22.5.

Table S9. Baseline serum bile acid concentration (μM) of all future metabolically healthy (MH) and unhealthy (MU) groups in the 10-year longitudinal study

|  | future metabolically healthy (MH) | future metabolically unhealthy (MU) |
| --- | --- | --- |
| HCA | 2.06E-02±2.60E-03 | 1.26E-02±1.10E-03* |
| HDCA | 1.04E-02±1.10E-03 | 4.67E-03±4.00E-04* |
| GHCA | 1.14E-02±1.66E-03 | 7.38E-03±9.20E-04* |
| GHDCA | 6.68E-02±6.36E-03 | 2.80E-02±1.88E-03* |
| NorDCA | 2.75E-03±5.50E-04 | 2.23E-03±3.90E-04 |
| 7-KetoLCA | 8.97E-03±1.23E-03 | 9.23E-03±9.90E-04 |
| UDCA | 9.17E-02±1.79E-02 | 9.84E-02±1.46E-02 |
| CDCA | 3.25E-01±5.55E-02 | 2.90E-01±3.83E-02 |
| DCA | 1.77E-01±2.44E-02 | 1.51E-01±1.53E-02 |
| 7-KetoDCA | 1.49E-02±7.70E-03 | 4.48E-03±7.20E-04 |
| 12-KetoCDCA | 4.70E-02±1.69E-02 | 2.38E-02±6.05E-03 |
| 3-ketoCA | 2.74E-03±7.10E-04 | 2.80E-03±8.80E-04 |
| CA | 1.31E-01±2.74E-02 | 1.17E-01±1.72E-02 |
| GLCA | 3.34E-03±9.00E-04 | 5.87E-03±1.58E-03 |
| GUDCA | 4.76E-02±6.04E-03 | 5.71E-02±8.14E-03 |
| GCDCA | 4.80E-01±5.08E-02 | 5.24E-01±5.56E-02 |
| GDCA | 8.31E-02±1.06E-02 | 9.24E-02±1.07E-02 |
| GCA | 1.68E-01±2.62E-02 | 1.73E-01±2.25E-02 |
| TLCA | 2.59E-03±8.70E-04 | 3.74E-03±2.05E-03 |
| TUDCA | 2.60E-03±5.70E-04 | 3.66E-03±6.80E-04 |
| TCDCA | 3.36E-02±8.12E-03 | 3.61E-02±4.45E-03 |
| TDCA | 1.45E-02±3.60E-03 | 1.63E-02±3.01E-03 |
| TCA | 1.63E-02±6.22E-03 | 1.68E-02±2.91E-03 |

Values were presented as mean ± S.E..

* p<0.05 comparing the two groups, Mann-Whitney test, after FDR correction.

Table S10. Baseline serum bile acid concentration (μM) of matched future metabolically healthy (MH) and unhealthy (MU) groups in the 10-year longitudinal study

|  | future metabolically healthy (MH) | future metabolically unhealthy (MU) |
| --- | --- | --- |
| HCA | 2.06E-02±2.57E-03 | 1.19E-02±1.49E-03* |
| HDCA | 1.04E-02±1.09E-03 | 4.55E-03±4.90E-04* |
| GHCA | 1.14E-02±1.64E-03 | 8.08E-03±1.46E-03* |
| GHDCA | 6.68E-02±6.29E-03 | 2.57E-02±2.36E-03* |
| NorDCA | 2.75E-03±5.50E-04 | 1.68E-03±4.30E-04* |
| 7-KetoLCA | 8.97E-03±1.21E-03 | 1.00E-02±1.54E-03 |
| UDCA | 9.17E-02±1.77E-02 | 1.08E-01±2.23E-02 |
| CDCA | 3.25E-01±5.49E-02 | 3.12E-01±5.41E-02 |
| DCA | 1.77E-01±2.41E-02 | 1.61E-01±2.41E-02 |
| 7-KetoDCA | 1.49E-02±7.62E-03 | 4.30E-03±9.00E-04 |
| 12-KetoCDCA | 4.70E-02±1.68E-02 | 1.79E-02±4.36E-03 |
| 3-ketoCA | 2.74E-03±7.00E-04 | 3.53E-03±1.61E-03 |
| CA | 1.31E-01±2.71E-02 | 1.14E-01±2.25E-02 |
| GLCA | 3.34E-03±8.90E-04 | 4.17E-03±1.24E-03 |
| GUDCA | 4.76E-02±5.98E-03 | 6.19E-02±1.29E-02 |
| GCDCA | 4.80E-01±5.03E-02 | 5.52E-01±8.44E-02 |
| GDCA | 8.31E-02±1.05E-02 | 8.44E-02±1.21E-02 |
| GCA | 1.68E-01±2.59E-02 | 1.78E-01±3.36E-02 |
| TLCA | 2.59E-03±8.60E-04 | 4.82E-03±3.69E-03 |
| TUDCA | 2.60E-03±5.60E-04 | 3.99E-03±9.90E-04 |
| TCDCA | 3.36E-02±8.03E-03 | 3.65E-02±6.24E-03 |
| TDCA | 1.45E-02±3.56E-03 | 1.52E-02±3.32E-03 |
| TCA | 1.63E-02±6.15E-03 | 1.81E-02±4.65E-03 |

Values were presented as mean ± S.E..

* p<0.05 comparing the two groups, Mann-Whitney test, after FDR correction.

Table S11. Metabolic markers of diabetic patients at baseline (0m) and 1, 3, 6, and 12 months after surgery in the gastric bypass surgery intervention study

|  | Baseline (0m) | 1 month after | 3 months after | 6 months after | 12 months after |
| --- | --- | --- | --- | --- | --- |
| n(M/F) | 38 (18/20) | 38 (18/20) | 38 (18/20) | 38 (18/20) | 38 (18/20) |
| Age (year) | 44.9±12.6 | / | / | / | 45.9±12.6 |
| BMI (kg/m2) | 32.2±3.8 | 28.1±3.5* | 26.0±3.2* | 24.8±2.9* | 24.5±2.7* |
| Waist (cm) | 107.2±12.5 | 95.9±9.8* | 89.9±9.8* | 86.6±9.1* | 86.1±8.5* |
| Glu0 (mmol/L) | 8.0±2.6 | 6.8±1.8* | 5.9±1.6* | 5.5±1.1* | 5.7±1.1* |
| Glu120 (mmol/L) | 12.3±4.0 | 7.5±2.5* | 7.4±3.0* | 6.9±3.0* | 6.8±2.7* |
| Ins0 (mU/L) | 25.5±22.4 | 12.9±10.4* | 8.0±5.0* | 7.1±4.4* | 7.3±4.2* |
| Ins120 (mU/L) | 105.2±79.1 | 21.0±17.6* | 23.8±20.7* | 31.9±32.5* | 24.2±19.8* |
| TC (mmol/L) | 7.7±1.9 | 6.7±1.2* | 6.0±1.2* | 5.9±0.6* | 5.9±1.2* |
| TG (mmol/L) | 2.6±3.1 | 1.5±0.6* | 1.2±0.6* | 1.1±0.6* | 1.0±0.6* |
| HDL (mmol/L) | 1.0±0.2 | 1.0±0.2* | 1.0±0.2 | 1.2±0.2* | 1.3±0.3* |
| LDL (mmol/L) | 3.1±1.0 | 3.2±1.0 | 2.6±0.6* | 2.5±0.8* | 2.5±0.6 |
| HbA1c (%) | 7.7±1.7 | 6.7±1.1* | 6.0±1.0* | 5.9±0.8* | 5.9±1.0* |
| HOMA-IR | 8.9±7.7* | 4.2±4.4* | 2.1±1.5* | 1.8±1.2* | 1.9±1.2* |

Values were presented as number or mean ± S.D..

* Wilcoxon paired samples signed-rank p<0.05 when compared with baseline (0m), after FDR correction.

BMI = body mass index; SP = [systolic](http://cn.bing.com/dict/clientsearch?mkt=zh-CN&setLang=zh&form=BDVEHC&q=收缩压) [blood](http://cn.bing.com/dict/clientsearch?mkt=zh-CN&setLang=zh&form=BDVEHC&q=收缩压) [pressure](http://cn.bing.com/dict/clientsearch?mkt=zh-CN&setLang=zh&form=BDVEHC&q=收缩压); DP = diastolic [blood](http://cn.bing.com/dict/clientsearch?mkt=zh-CN&setLang=zh&form=BDVEHC&q=收缩压) [pressure](http://cn.bing.com/dict/clientsearch?mkt=zh-CN&setLang=zh&form=BDVEHC&q=收缩压); Glu0 = fasting plasma glucose; Glu120 = 2 h plasma glucose; Ins0 = fasting insulin; Ins120 = 2 h insulin; TG = total triglycerides; TC = total cholesterol; HDL = high density lipoprotein; LDL = low density lipoprotein; HOMA-IR = Glu0×Ins0/22.5.

Table S12. Serum bile acid concentration (μM) of diabetic individuals at baseline (0m) and 1, 3, 6, and 12 months after gastric bypass surgery

|  | Baseline (0m) | 1 month after | 3 month after | 6 month after | 12 months after |
| --- | --- | --- | --- | --- | --- |
| HCA | 7.49E-03±1.64E-03 | 2.07E-02±2.22E-03* | 2.12E-02±2.23E-03* | 2.23E-02±2.22E-03* | 2.33E-02±2.22E-03* |
| HDCA | 3.86E-03±1.09E-03 | 1.49E-02±1.08E-03* | 1.48E-02±1.08E-03* | 1.59E-02±1.15E-03* | 1.56E-02±1.18E-03* |
| GHCA | 6.20E-03±1.56E-03 | 1.42E-02±1.56E-03* | 1.54E-02±1.56E-03* | 1.58E-02±1.57E-03* | 1.61E-02±2.45E-03* |
| GHDCA | 2.25E-02±5.57E-03 | 5.22E-02±5.54E-03* | 5.59E-02±5.74E-03* | 6.16E-02±5.63E-03* | 6.12E-02±5.62E-03* |
| NorDCA | 1.52E-03±1.70E-04 | 2.93E-03±1.70E-04* | 2.69E-03±1.70E-04* | 2.46E-03±1.80E-04 | 2.67E-03±1.80E-04 |
| 7-KetoLCA | 1.80E-02±3.17E-03 | 9.47E-03±3.03E-03* | 1.56E-02±3.06E-03 | 9.65E-03±3.06E-03* | 1.15E-02±3.06E-03* |
| UDCA | 1.19E-01±2.07E-02 | 5.88E-02±1.81E-02* | 8.31E-02±1.82E-02 | 9.02E-02±1.81E-02* | 8.93E-02±1.81E-02 |
| CDCA | 3.39E-01±4.25E-02 | 1.95E-01±4.12E-02* | 1.65E-01±4.05E-02* | 1.69E-01±4.07E-02* | 1.86E-01±4.06E-02* |
| DCA | 1.46E-01±2.31E-02 | 9.58E-02±2.33E-02 | 1.61E-01±2.33E-02 | 1.83E-01±2.32E-02 | 2.07E-01±2.32E-02 |
| 7-KetoDCA | 9.79E-03±2.38E-03 | 6.38E-03±2.38E-03 | 2.64E-02±2.38E-03* | 7.22E-03±2.37E-03 | 1.15E-02±2.37E-03 |
| 12-KetoCDCA | 3.18E-02±1.69E-02 | 3.80E-02±1.68E-02* | 3.46E-02±1.68E-02 | 2.69E-02±4.50E-03 | 2.52E-02±4.52E-03 |
| 3-ketoCA | 2.12E-02±2.73E-03 | 2.75E-02±2.81E-03 | 5.51E-02±2.81E-03 | 1.70E-02±2.77E-03 | 3.34E-02±2.82E-03 |
| CA | 7.58E-02±2.11E-02 | 8.64E-02±2.65E-02 | 2.47E-01±2.65E-02 | 1.04E-01±2.65E-02 | 1.36E-01±2.65E-02 |
| GLCA | 5.91E-03±1.78E-03 | 5.01E-03±1.78E-03 | 5.60E-03±1.78E-03 | 7.31E-03±1.78E-03 | 9.75E-03±1.78E-03* |
| GUDCA | 4.07E-02±1.03E-02 | 3.50E-02±9.57E-03 | 2.33E-02±9.57E-03* | 3.17E-02±9.57E-03* | 2.41E-02±1.10E-02* |
| GCDCA | 4.35E-01±6.36E-02 | 5.31E-01±6.37E-02 | 5.08E-01±6.43E-02 | 6.42E-01±7.33E-02* | 6.63E-01±7.42E-02* |
| GDCA | 7.13E-02±2.20E-02 | 5.04E-02±2.21E-02 | 6.01E-02±2.20E-02 | 8.26E-02±2.20E-02 | 6.53E-02±2.24E-02 |
| GCA | 1.66E-01±3.17E-02 | 2.25E-01±3.18E-02 | 2.11E-01±3.19E-02 | 2.61E-01±3.19E-02* | 1.74E-01±3.19E-02 |
| TLCA | 3.55E-03±9.00E-04 | 3.26E-03±9.00E-04 | 2.48E-03±9.00E-04 | 2.79E-03±9.00E-04 | 3.59E-03±9.00E-04 |
| TUDCA | 4.89E-03±6.90E-04 | 9.93E-03±6.80E-04 | 4.50E-03±6.90E-04 | 5.75E-03±7.00E-04 | 3.32E-03±8.20E-04* |
| TCDCA | 4.22E-02±9.80E-03 | 7.72E-02±9.81E-03* | 7.46E-02±9.83E-03 | 5.77E-02±9.84E-03 | 4.84E-02±1.26E-02 |
| TDCA | 1.47E-02±4.43E-03 | 1.96E-02±4.43E-03 | 1.96E-02±4.41E-03 | 1.92E-02±4.42E-03 | 1.55E-02±4.52E-03 |
| TCA | 1.67E-02±4.11E-03 | 3.32E-02±4.10E-03 | 2.96E-02±4.09E-03 | 2.32E-02±4.09E-03 | 1.80E-02±5.26E-03 |

Values were presented as mean ± S.E..

* p<0.05 when compared with Baseline (0m), Wilcoxon paired samples signed-rank test, after FDR correction.

**Human experiments**

The four human studies reported in this paper were all approved by the Ethics Committee of Shanghai Jiao Tong University Affiliated Sixth People's Hospital in accordance with the World Medical Association’s Declaration of Helsinki. Written informed consent was obtained from all participants before recruitment.

**Cell Studies**

*Cell culture and Reagents*

Human cecum cell line NCI-H716 (CCL-251) and mouse colon cell line STC-1 (CRL-3254) were purchased from ATCC. The cells were maintained at 37 ^o^C, 5% CO_2_, in RPMI 1640 medium (Invitrogen) supplemented with 10% fetal bovine serum (FBS, Gibco, USA). shRNA plasmids targeting human and mouse TGR5 (sc-61678-SH for human, sc-61679-SH for mouse) and FXR (sc-38848-SH for human and sc-155894-SH) were purchased from SANTA CRUZ Biotech. The cells were transfected by various shRNA plasmids and screened for stable transfection using puromycin. The knockdown of target proteins was verified by western-blot. The cell line was tested for mycoplasma contamination, and treated with different BAs, BAs + 5β-CA, or BAs + CDCA, as the experiments described following.

1) NCI-H716 and STC-1 cells were plated into 24 well-plates with an initial cell density of 5 ×10^5^ cells/well, treated with different concentrations (5, 25, and 50 μM) of different BAs, including 6 HCA species (HCA, THCA, GHCA, HDCA, THDCA, and GHDCA), and 19 other BAs (CA, CDCA, LCA, DCA, UDCA, αMCA, βMCA, ωMCA, TCA, GCA, TCDCA, GCDCA, TDCA, GDCA, TLCA, GLCA, TαMCA, TβMCA, and TωMCA) with 48-hour treatment. The proglucagon GCG and secreted GLP-1 of all the cells were determined by qRT-PCR and ELISA kits, respectively. The intracellular GLP-1 of the cells treated with 25μM of BAs including 6 HCA species (HCA, THCA, GHCA, HDCA, THDCA, and GHDCA), and 8 other representative BAs (CA, CDCA, LCA, DCA, UDCA, αMCA, βMCA, ωMCA) were determined by western blot.

2) The expression of TGR5 was knocked down in both NCI-H716 and STC-1 cells. NCI-H716 and STC-1 control and knockdown cells were treated with 6 HCA species (HCA, THCA, GHCA, HDCA, THDCA, and GHDCA) at 25 μM for 48h. The intracellular GLP-1, CREB phosphorylation, CREB, and α-Tubulin of the cells were determined by western blot. The secreted GLP-1 was determined by ELISA kits.

3) NCI-H716 cells were treated with different BAs, including 6 HCA species (HCA, THCA, GHCA, HDCA, THDCA, and GHDCA), and other representative BAs (CA, GCA, CDCA, GCDCA, LCA, and GLCA) at 25 and 50 μM with 48-hours of treatment. The nuclear and cytoplasmic FXR, LAMIN A, β-ACTIN, SHP, P-CREB, and CREB of the cells were determined by western blot.

4) The NCI-H716 cells were treated with CDCA or 5β-CA at 50 μM for 24 hours, with or without the presence of 50 μM HCA species (HCA, THCA, GHCA, HDCA, THDCA, and GHDCA). The nuclear and cytoplasmic FXR, LAMIN A, and β-ACTIN of the cells were determined by western blot.

5) The NCI-H716 cells were treated with 6 HCA species (HCA, THCA, GHCA, HDCA, THDCA, and GHDCA), 6 HCA species with 5β-CA and other representative BAs (TCA, TCDCA, TLCA, TDCA, TUDCA, GCA, GCDCA, GLCA, GUDCA) at 50 μM for 48 hours. The proglucagon GCG and secreted GLP-1 of all the cells were determined by qRT-PCR and ELISA kits, respectively. The intracellular GLP-1 and α-ACTIN of the cells with HCAs with or without 5β-CA were determined by western blot and immunofluorescence staining.

6) The expression of FXR was knocked down in NCI-H716 cells. NCI-H716 control and knockdown cells were treated with 6 HCA species (HCA, THCA, GHCA, HDCA, THDCA, and GHDCA) and other representative BAs (CA, CDCA, LCA, and DCA) at 25 and 50 μM for 48h. The proglucagon GCG of all the cells were determined by qRT-PCR, and secreted GLP-1 of the cells treated with 50 μM BAs were determined by ELISA kits. The intracellular TGR5, pCREB, CREB, and β-ACTIN of the cells treated with 50μM BAs were determined by western blot.

*RNA isolation and quantitative reverse transcription PCR (qRT-PCR)*

Total RNA was isolated using Trizol (Thermo Fisher Scientific, CA) according to the manufacturer’s instruction. The cDNA was synthesized from the total RNA using iScript Reverse Transcription Supermix for RT-qPCR (Bio-Rad CA). Transcript levels were measured in triplicate by qRT-PCR (LightCycler 480 II, Roche) using iTaq™ Universal Probes Supermix (Bio-Rad CA), with the Taqman probes targeting proglucagon GCG (Ref. #162836223, IDT) and beta-actin ACTB (Ref. #162655388, IDT).

*Immunofluorescence Staining for intracellular GLP-1*

The cells were stained with antiGLP-1 antibody (ab23472, Abcam, Cambridge, MA), which binds to both precursor and GLP-1 cleavage products. For IF staining of mouse Ileum tissues, the tissues were embedded with Tissue-Tek O.C.T. Compound (Sakura Finetek U.S.A., Inc. CA), the slides were staining with GLP-1(ab23472), TGR5(ab72608), p-CREB S133(ab220798) and FXR (ab28480)(All purchased from Abcam, Cambridge, MA). The slides were then stained with Goat Anti-Rabbit IgG H&L (FITC) (1:1000, ab6717), and Goat Anti-Mouse IgG H&L (Alexa Fluor® 594) (1:1000, ab150116) (Cambridge, MA). The slides were mounted using ProLong™ Gold Antifade Mountant (P36930, ThermoFisher), and were photographed using a digitalized microscope camera (Nikon, Tokyo, Japan). Positive cells were quantified for Mean Florescence Density (MFI) using Image J software (NIH).

*Western blot	for intracellular GLP-1*

The whole cell lysates were prepared using RIPA Lysis and Extraction Buffer (89900, Thermo Scientific). The nuclear and cytoplasm fractions were separated using NE-PER™ Nuclear and Cytoplasmic Extraction Reagents (78833, Thermo Scientific). Equal amounts of proteins were boiled with 4×laemmli buffer (Catalog #161-0747, Bio-rad), separated on 10% SDS-PAGE, transferred onto a PVDF membrane, probed by antibodies for GLP-1 (ab23472, Abcam), TGR5 (ab72608, Abcam), FXR (WH0009971M1, Sigma),CREB (9104, Cell Signaling),	Phospho-CREB-Ser133 (9198, Cell Signaling) and β-actin (ab6276, Abcam), and visualized via an ECL kit (170-5060, Bio-rad). Antibodies to GLP-1 (ab23472), β-actin (ab6276) and α-Tubulin (ab18251) were purchased from Abcam (Cambridge, MA).

*ELISA for Secreted GLP-1*

The cell media was removed from the cells, centrifuged at 2,000 rpm for 10 min to remove cell debris and the supernatant was collected. GLP1 concentration was determined using a Human GLP-1 ELISA Kit according to the manufacturer’s instruction (ab184857, Abcam, MA).

**Quantitative analysis of BAs**

BAs were quantified using our established methods with minor modifications to improve accuracy.

*Sample pretreatment for BA analysis*

An aliquot of 25 µL serum sample was mixed with 150 µL acetonitrile-methanol (8:2 v/v) containing 6 internal standards (IS) (D4-GCA, D4-GDCA, D4-CA, D4-UDCA, D4-LCA, and D4-GCDCA, 50 nM for each). The mixture was allowed to stand at 20 °C for 30 min, and was then centrifuged at 13,000 rpm at 4 °C for 30 min. An aliquot of 150 µL of supernatant was transferred to another tube and then vacuum-dried. A 25 µL acetonitrile-methanol (8:2 v/v) were added, the sample was re-vortexed at 1,500 rpm, 10 °C for 10 min and 25 µl water containing 0.0% formic acid were added. The sample was vortexed again at 1,500 rpm, 10 ^o^C for 10 min, and then centrifuged at 13,000 rpm, 4 ^o^C for 15 min. The supernatant was used for UPLC-MS analysis.

*Instrument analysis*

An aliquot of 5 µM standard stock solution was prepared by mixing BA standards. A series of standard calibration solutions were diluted with desalted serum (depleted of BAs using activated charcoal) for the calibration curve. The calibration curve and the corresponding regression coefficients were obtained by internal standard adjustment. A Waters ACQUITY ultra performance LC system equipped with a binary solvent delivery manager and a sample manager (Waters, Milford, MA) was used throughout the study. The mass spectrometer was a Waters XEVO TQ instrument with an ESI source (Waters Corp., Milford, MA). The entire LC−MS system was controlled by MassLynx 4.1 software. All chromatographic separations were performed with an ACQUITY BEH C18 column (1.7 μm, 100 mm × 2.1 mm internal dimensions) (Waters Corp., Milford, MA). The mobile phase consisted of water with 0.01% formic acid (mobile phase A) and acetonitrile/methanol (87/13, v/v, mobile phase B). The flow rate was 0.45 mL/min with the following mobile phase gradient: 0−0.5 min (5% B), 0.5−1 min (5−20%B), 1−2 min (20−25%B), 2−5.5 min (25% B), 5.5−6 min (25−30%B), 6−8 min (30% B), and 8−9 min (30−35% B), 9−17 min (35−65%B), 17−18 min (65−99%B), 18−19 min (99%B), 19−19.1 min (99−5%B), 19−20 min (5%B). The column was maintained at 45 ^o^C and the injection volume for all samples was 5 μL. The mass spectrometer was operated in negative ion mode with a 2.5 kV capillary voltage. The source and desolvation gas temperature were 150 and 450 ^o^C, respectively. The data were collected with a multiple reaction monitor (MRM), and the cone and collision energy for each BA used the optimized settings from QuanOptimize application manager (Waters Corp., Milford, MA).
